# Profiling the endothelial translatome *in vivo* using ‘AngioTag’ zebrafish

**DOI:** 10.1101/815696

**Authors:** Mayumi F. Miller, Leah J. Greenspan, Derek E. Gildea, Kathryn Monzo, Gennady Margolin, Van N. Pham, Keith Ameyaw, Lisa Price, Natalie Aloi, Amber N. Stratman, Andrew E. Davis, Isabella Cisneros, Caleb Mertus, Ryan K. Dale, Andreas D. Baxevanis, Brant M. Weinstein

## Abstract

Vascular endothelial cells *in vivo* are exquisitely regulated by their local environment, which is disrupted or absent when using methods such as FACS sorting of cells isolated from animals or *in vitro* cell culture. Here, we profile the gene expression patterns of undisturbed endothelial cells in living animals using a novel “AngioTag” zebrafish transgenic line that permits isolation of actively translating mRNAs from endothelial cells in their native environment. This transgenic line uses the endothelial cell-specific *kdrl* promoter to drive expression of an epitope tagged Rpl10a 60S ribosomal subunit protein, allowing for Translating Ribosome Affinity Purification (TRAP) of actively translating endothelial cell mRNAs. By performing TRAP-RNAseq on AngioTag animals, we demonstrate strong enrichment of endothelial specific genes and uncover novel endothelial genes and unique endothelial gene expression signatures for different adult organs. Finally, we generated a versatile “UAS:RiboTag” transgenic line to allow a wider array of different zebrafish cell and tissue types to be examined using TRAP-RNAseq methods. These new tools offer an unparalleled resource to study the molecular identity of cells in their normal *in vivo* context.

## INTRODUCTION

The vascular system is a complex network of arteries, veins, and capillaries working in concert to allow oxygenated blood to flow throughout the body and transport hormones and small molecules to target tissues. This network is comprised of lumenized endothelial cell tubes surrounded by smooth muscle cells (in larger vessels, especially arteries) or pericytes juxtaposed to the basement membrane. Endothelial cells in vascular tubes have extensive cell-cell junctional contacts with one another and with the surrounding vascular smooth muscle cells. In addition, the luminal surface of the endothelium is constantly exposed to flow dynamics, circulating cells, and circulating factors from the blood, while the abluminal surface interacts extensively with extracellular matrix and with numerous non-vascular cell types and tissues. Understanding how endothelial cells respond to this complex multitude of external cues and signals *in vivo* to regulate blood vessel growth and function is of enormous interest given the important roles vessels play in a variety of pathologies including cancer, ischemia, and congenital vascular disorders.

Existing methods for analysis of endothelial gene expression and function generally rely on post-mortem single-cell isolation and/or *in vitro* culture of endothelial cells, disrupting virtually all of the normal interactions experienced by endothelial cells *in vivo.* A number of animal models have been used to study endothelial cells in their endogenous setting, including mice, birds, and zebrafish. The zebrafish is a particularly useful model organism for studying the vasculature, with externally developing, optically clear embryos and larvae, and a variety of transgenic reporter lines expressing fluorescent proteins in the endothelium that permit high-resolution visualization of the complex and widely dispersed vascular network at all stages of development including adults^1,2^. These features, along with the ability to house large numbers of adult animals and obtain large numbers of progeny, facilitate genetic and experimental analysis of vascular development and homeostasis in the fish. The advantages of the zebrafish model have led to important discoveries including mechanisms of arterial-venous differentiation^3–6^, lumen formation^7–9^, and guidance and patterning of vessel networks during development^10,11^.

Although zebrafish have been invaluable for observing endothelial cell behavior *in vivo,* existing methods for global analysis of endothelial gene expression such RNAseq or single-cell RNAseq still require enzymatic and mechanical dissociation into single-cell suspensions that are then subjected to further often lengthy sorting procedures before RNA is prepared from the cells. It is likely that these manipulations introduce signaling pathway changes, including activation of cell stress and apoptotic pathways, that obscure the native transcriptional state of endothelial cells. In contrast, Translating Ribosome Affinity Purification (TRAP)^12^ involves collecting actively translating mRNAs by immunoprecipitating the attached ribosomes translating these mRNAs. Studies in mice have demonstrated that transgene-driven expression of an epitope-tagged ribosomal protein subunit (also known as a “RiboTag”) within specific cell types or tissues can be used to affinity purify and profile actively translating mRNAs *in vivo* by performing TRAP followed by RNAseq (TRAP-RNAseq) or microarray experiments^12–14^.

We have now developed transgenic zebrafish that permit profiling of global gene expression in vascular endothelial cells (“AngioTag”) or in any cell or tissue type for which a Gal4 driver line is available (“UAS:RiboTag”). These lines utilize the vascular endothelial kdrl or UAS promoters, respectively, to drive co-expression of HA-tagged Rpl10a (a 60S ribosomal subunit protein) and EGFP (a visible transgene marker), permitting immunoprecipitation of tissue-specific ribosomes and collection of the mRNAs they are attached to and translating. Since isolation of vascular mRNA from these transgenic lines is extremely rapid (embryos and organs are quickly homogenized in a lysis buffer containing cycloheximide to halt further translation), the mRNAs collected represent the *bona fide* expression profiles of undisrupted cells *in situ.* Using this tool, we have uncovered novel endothelial genes involved in embryonic vessel development as well as unique endothelial signatures of vessels in different adult organs. These new transgenic lines provide powerful tools for profiling cell- or tissue-specific gene expression in the zebrafish that can be used to study changes in gene expression during development, disease, or repair after injury.

## METHODS

### Zebrafish

Zebrafish were maintained and zebrafish experiments were performed according to standard protocols^15^ and in conformity with the Guide for the Care and Use of Laboratory Animals of the National Institutes of Health, in an Association for Assessment and Accreditation of Laboratory Animal Care (AAALAC) accredited facility. Fish were housed in a large recirculating aquaculture facility wih 1.8L and 6L tanks. Water quality was routinely measured and proper parameters taken to maintan water quality stability. Fry were fed rotifers and adults were fed Gemma Micro 300 (Skretting). The following transgenic fish lines were used in this study: EK (wild-type), *Tg(kdrl:gfp)^la^*^116^ ^16^, *Tg(kdrl:egfp-2a-rpl10a3xHA)^530^*^530^ (this paper), *Tg(uas:egfp-2a-rpl10a2xHA)^530^*^531^ (this paper), *Tg(xa210:gal4)^530^*^241^ ^17^, *Tg(fli1a:gal4ff)^ubs^*^4^ ^18^, and *Tg(huc:gal4)* ^19^.

### Sucrose Gradient

Sucrose density gradients were prepared for sedimentation analysis of polysome profiles. 12 mL 5-50% sucrose gradients were prepared in 110 mM KOAc, 2M MgOAc, 10 mM HEPES pH 7.6 with BioComp Gradient Master (BioComp), and allowed to rest overnight at 4°C. Dechorionated and deyolked embryos were dounce homogenized in 1.5 µl polysome fractionation buffer per embryo (10mM HEPES pH7.4, 110mM KOAc, 2mM MgOAc, 100mM KCl, 10mM MgCl_2_, 0.1% Nonidet P-40, 2mM DTT, 40U/mL RNasin, 200ug/mL cycloheximide, and protease inhibitors; adapted from^20^). Homogenates were centrifuged at 1000xg for 10 min at 4°C. Protein concentration was quantified by Bradford Assay (Sigma Aldrich) and equivalent amounts were loaded onto gradients. Samples were centrifuged in a SW41 Beckman rotor at 40000 rpm at 4°C for 2 hours. 16 x 1mL fractions were collected with an ISCO piercing apparatus connected to a BioLogic chromatography system at 0.5 mL/min, pushing 55% sucrose. Data were collected using LP DataView software (BioRad). For EDTA treatment, samples were treated with 200 mM final concentration EDTA prior to loading onto gradient.

### TRAP protocol

Translating Ribosome Affinity Purification (TRAP) was performed as described previously^12^ with modifications. For each larval TRAP sample approximately 1200-1500 24hpf zebrafish embryos were dechorionated, deyolked, and dounce homogenized in homogenization buffer consisting of 50 mM Tris pH 7.4, 100 mM KCl, 12 mM MgCl_2_, 1% NP-40, 1 mM DTT (Sigma, Cat. #646563), 1x Protease inhibitors (Sigma, Cat. #P8340), 200 units/mL RNAsin (Promega, Cat. #N2115), 100ug/mL Cycloheximide (Sigma Cat. #7698), 1mg/mL Heparin (Sigma, Cat. #H3393-10KU). Lysates were cleared for 10 min at 10,000xg, 4°C. 1 µl of anti-HA antibody (Abcam ab9110 Rabbit polyclonal) was added per 400µg protein in an 800 µl lysate and the samples were orbitally rotated at 4°C for 5 hours. 60 µl of Dynabeads® protein G slurry (Novex/Life Technologies) was added per 1 µl of anti-HA antibody with homogenization. Before using, Dynabeads were washed with 800ul homogenization buffer, rotating for 30 minutes to equilibrate the beads. The wash buffer was then removed from the Dynabeads and the lysate + Ab solution was added to the equilibrated Dynabeads. This Dynabead + lysate + Ab mixture was incubated overnight at 4°C on an orbital rotator. Dynabeads were collected using a magnetic stand, and washed 3 x 5min on an orbital shaker with high salt homogenization buffer (50mM Tris pH 7.4, 300mM KCl, 12mM MgCl_2_, 1% NP-40, 1mM DTT, 1x Protease inhibitors, 200 units/mL RNAsin, 100µg/mL Cyclohexamide, 1mg/mL Heparin). RNA was collected from the Dynabeads and DNAse treated using the Zymo ZR-Duet Kit (Zymo Research D7003). Note that careful attention to the protocol is required to keep lysates cold and free of RNases, and RNA integrity must be carefully monitored. Large numbers of embryos are needed to provide the starting material for the TRAP protocol because (i) the endothelial or other cell types expressing the RiboTag represent a small fraction of the total cells in the animal, (ii) only a fraction of the ribosomes in the targeted cell type contain the RiboTag - most ribosomes are still “untagged,” and (iii) TRAP-purified RNA samples contain a large amount of co-purifying ribosomal RNAs in addition to the mRNAs, and (iv) we prepared largely unamplified libraries for sequencing.

A similar TRAP protocol was utilized for the adult whole fish and organ samples as described above, with a few additional modifications. SUPERase-In (100U/µl) was added to the homogenization buffer as an additional RNAse inhibitor. Organs were either dissected fresh or flash frozen for later use. Whole fish were flash frozen and ground into fish powder before homogenization. Four organs/whole fish were used per replicate and the amount of each tissue was weighed so only 0.75% - 3% tissue to homogenization buffer (weight/volume) was utilized. Tissue was homogenized using a Cole-Parmer LabGen 850 homogenizer at 13,000rpm for 45s. 5µl of anti-HA antibody was used per sample. Antibody and Dynabeads incubation was the same as the larval samples but RNA was collected from the Dynabeads using RNeasy Micro Plus Kit (Qiagen #74034) with RNA-only ß-Mercaptoethanol supplementation in the RLT buffer.

### Quantitative RT-PCR

RNA was reverse transcribed using a High Capacity cDNA reverse transcription kit (Applied Biosystems™). cDNA was combined with the primers listed below and with LightCycler® 480 SYBR Green I (Roche). Reactions were run in a LightCycler® 480 Multiwell Plate 384 machine (Roche). Relative fold changes were calculated using ddCT calculations and by calculating standard deviations. The following primer pairs were used:

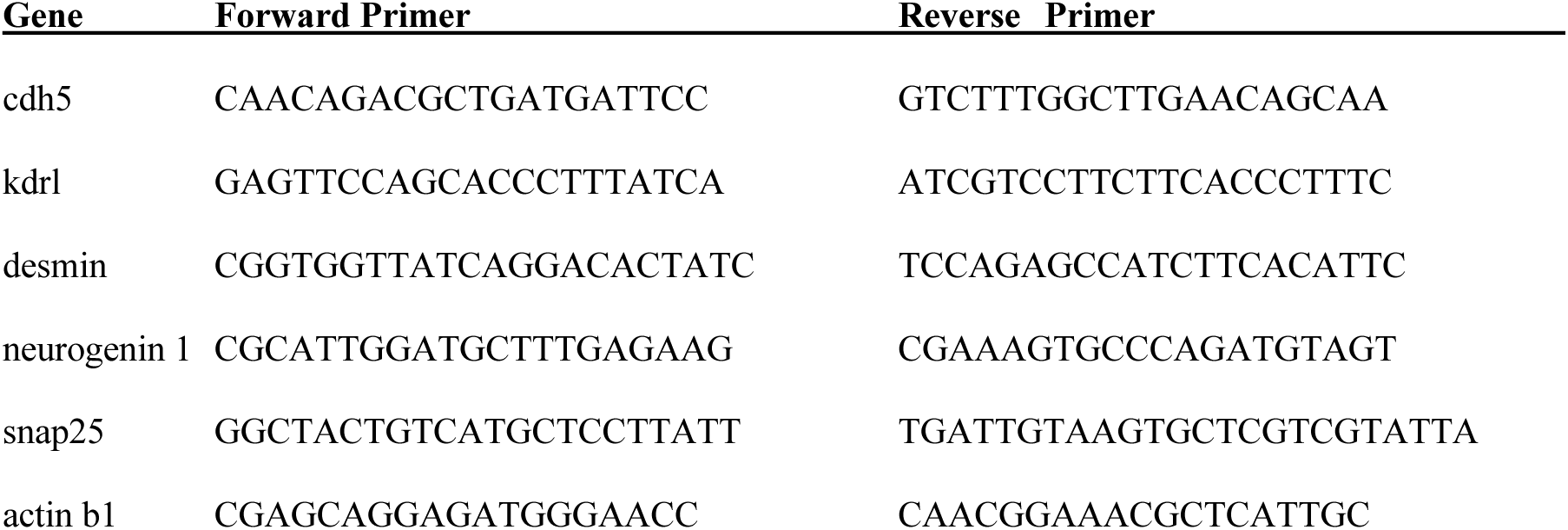

### RNAseq

RNA concentration and quality were measured on a Qubit (Thermo Fisher) and on an Agilent 2100 BioAnalyzer + RNA Pico Chip (Agilent), respectively. For our 24hpf TRAP samples, a large proportion of the RNA samples we collected represented ribosomal RNAs co-purified with our polysome mRNAs, so 500 ng RNA was polyA selected or rRNA depleted using Ribo-Zero rRNA Removal Kit (Illumina) before library construction. Sequencing libraries were constructed from the purified mRNA using TruSeq Stranded mRNA Library Prep Kits (Illumina). Libraries were sequenced using an Illumina HiSeq 2500 to generate approximately 50 million 2 x 75bp or 2 x 100bp paired end reads. Raw data were de-multiplexed and analyzed. RNAseq data was deposited with GEO, accession # GSE121973.

For adult whole fish and organs, RNA samples were submitted to Novogene, Inc. for their ultralow input mRNA sequencing. Samples underwent quality control, library construction, and sequencing. RNA libraries for RNAseq were prepared by Novogene using SMART-seq V4 Ultra Low Input RNA kit & non-directional library. Messenger RNA was enriched using a poly-T oligo-attached magnetic beads. Samples were sequenced on a Illumina NovaSeq 6000 to generate 150bp pair end reads ranging from 24 - 94 million raw reads per sample. RNAseq data was deposited with GEO, accession # GSE276280.

### Assessing mRNA abundance from RNAseq data

For our larval TRAP samples, RNAseq reads were mapped to the zebrafish reference genome (GRCz10) using the STAR alignment software package (v. 2.4.2a). Only those RNAseq reads mapping to annotated protein-coding Ensembl gene models (release 85) were used for gene expression profiling. Quantification of the RNAseq data was done using the RSEM software suite (v. 1.2.22), obtaining expected and normalized RNAseq read counts (in transcripts per million, TPM). DESeq2 was used to compare mRNA abundance between samples. For comparing mRNA abundance between TRAP-enriched and FACS-sorted samples, expected RNAseq read counts from enriched samples were normalized to the read counts from RNAseq performed on total mRNA input used for the TRAP enrichment and total mRNA from unsorted cells prior to FACS. Using the RNAseq data from the TRAP input and the unsorted cells prior to FACS, normalization factors were calculated by dividing the TPM abundance for a given mRNA in each sample by the geometric mean of the TPM abundances across all samples. The expected read counts from the TRAP-RNAseq and FACS-RNAseq data sets were divided by their corresponding normalization factor to adjust for differences in input mRNA levels prior to enrichment. With these adjusted RNAseq read counts, DESeq2 was used to identify differences in mRNA abundance.

For the adult TRAP samples, sequenced FASTQ files were trimmed with cutadapt and aligned with hisat2 using lcdb-wf pipeline (https://github.com/lcdb/lcdb-wf, v1.9rc). We used zebrafish genome assembly version GRCz11 and Ensembl gene annotation version 99. Subsequent downstream analyses were performed with the lcdb-wf pipeline utilizing R and packages DESeq2, clusterProfiler and GeneTonic (for fuzzy clustering). To determine which genes were endothelial enriched within a specific tissue, one-sided Wald t-tests were used to select for genes that were upregulated in the TRAP pulldown (IP) for that tissue compared to its total mRNA lysate (input), later refered to as the IP>Input comparison. To determine which endothelial enriched genes were shared among all tissues and the whole fish control, a multiple testing procedure for multi-dimensional pairwise comparisons was applied^21^. To control the false discovery rate (FDR) across genes at 10%, we extracted raw p-values for each gene and each tissue IP > Input comparison, with missing values being replaced by 1. A Holm correction was applied across comparisons for each gene separately. The FDR was calculated across genes using gene-wise minimal Holm-corrected p-values, and the fraction R of genes with FDR < 0.1 was determined. For each IP > Input comparison we selected genes whose within-gene Holm-corrected p-values were less than 0.1*R (and whose across-genes FDR was less than 0.1). Intersection of these genes across all IP > Input comparisons yielded a set of 350 genes. Out of this list, select genes were picked to be displayed in a heatmap in which their TRAP pulldown log2fold expression for each tissue was ≥ 1.2 compared to their own tissue lysate with a p-adjusted value <0.05.

To determine which genes were uniquely enriched in the endothelium of specific organs, we pre-screened for genes within each tissue whose TRAP pulldown (IP) was upregulated compared to its total tissue mRNA lysate (input) and, separately, whose TRAP pulldown was upregulated compared to the whole fish TRAP pulldown, using a similar multiple testing procedure for multi-dimensional pairwise comparisons as described above, and taking an intersection of the two comparions. This yielded a list of 3304 genes combined across all tissues (excluding whole fish). Based on this gene set, we determined genes dominantly expressed in each tissue’s TRAP pulldown (IP) sample. The criteria for selection were as follows: 1) a gene was selected in a given tissue during the pre-screening, 2) it passed the omnibus FDR < 0.1 testing overall gene change across IPs (using minimum of Holm-corrected p-values of all pairwise IP versus IP comparisons), 3) statistically verified greater IP expression than in all other tissue IPs (with cross-IP Holm-corrected p-value < 0.1*R, where R is the fraction of genes passing the omnibus test), 4) the minimum across the tissue IP replicates is above any other IP replicates in all other tissues. Gene lists for each tissue were then further filtered as follows: 1) endothelial gene expression was at least 1.5-fold greater than that of the organ lysate, 2) endothelial gene expression was at least 2-fold greater than the endothelial gene expression of the next highest tissue, 3) the difference in endothelial gene expression between the tissue of interest and the next highest expressing tissue was at least 2-fold greater than their tissue lysate differences. A heatmap was generated using the top 7-8 genes that fit these criteria for each organ.

### Analysis of gene features

Ensembl gene models were used to calculate codon usage and 3’ UTR lengths. The longest transcript/isoform was selected to represent each annotated Ensembl gene. For calculating codon usage, the coding sequences (CDS) for all transcripts were obtained from Ensembl (release 87, ftp://ftp.ensembl.org) in FASTA format. Custom Perl scripts were used to retrieve specific transcript sequences and to count codons from these sequences. For calculating 3’ UTR lengths, genomic position information for Ensembl transcripts was downloaded from UCSC Table Browser in genePred text format. The genomic start position for the 3’ UTR for the longest transcript of each Ensembl-annotated gene was obtained using the nucleotide position immediately following the end of the annotated coding sequence position. The 3’ UTR stop position was obtained from the annotated transcription stop site. The 3’ UTR length was calculated using these start and stop positions.

### Western blot

Protein samples were collected and boiled in Laemmli buffer + 2-Mercaptoethanol and run on 12% polyacrylamide gels, transferred onto PVDF membranes, and then blocked in 5% BSA. Blots were probed with either anti-RPL11 (Abcam, Cat. # ab79352) or anti-HA (Sigma, H9658) antibodies. Blots were exposed on film (Amersham Hyperfilm ECL, 28906836) using Amersham ECL Western Blotting Reagent (GE, RPN2106).

### Fluorescence Activated Cell sorting

For Fluorescence Activated Cell Sorting (FACS), 24hpf AngioTag embryos were anaesthetized with Tricaine and dechorionated and deyolked. Embryos were then washed in PBS and incubated in 0.25% trypsin at room temperature with trituration until dissociated into a single cell suspension. Following embryo dissociation, trypsin was quenched with Leibovitz L-15 phenol-free media with 10% FBS (Gibco). Cells were passed through a 70 uM strainer and collected by centrifugation at 600xg for 1 min. Cells were resuspended in phenol-free L-15 + 10% FBS, and GFP+ cells were collected by FACS sorting on a BD FACSAria™ III machine using BD FACSDiva™ software (Becton Dickinson Biosciences). Sorted cells were collected at 500xg for 5 min and RNA was isolated and DNAse treated using the ZR-Duet kit (Zymo Research D7003).

### Whole-mount RNA *in situ* hybridization

DIG-labeled antisense riboprobes for the below genes were generated using DIG Labeling Kit (Roche). *In situ* hybridization was performed as described^3^. BM purple (Sigma) was used for DIG-labeled probes. Riboprobes were generated for the following genes:

ENSDARG00000069998 (F’ GGTGCAGATAACTGGGAAGGTGATAG,

R’ TCAGTGTGAAGACGTACACC),

ENSDARG00000076721 (F’ CAGATGAAGTAAAGTCAGTATCTGTGATG,

R’ GTAGTCTGGTTGGTGAATGAATAAGC),

ENSDARG00000098293 (F’ GACCATGTGCTGAGAAATGTGAAGAG,

R’ TGTTAGCTCCATTTCCGCAG),

ENSDARG00000098129 (F’ GCTGTCTGTGGAGCGCTAAGTGTTTGTCT,

R’ TAATACGACTCACTATAGGGAGAATGTCACATCCGACCAATCAGAAT),

ENSDARG00000093124/scpp8 (F’ GATGAATACTTTGAAGGGATTGATTCTGAT, R’

CATCATAGCATCAGAAATCATCAAG),

ENSDARG00000008414/exoc312a (F’ GAGGCTGAAGGTAGATTTGGACAGATCGAC,

R’ TGTGGATCCCACTATTTTTACAGTG),

ENSDARG00000056643/slc22a7b.1 (F’ ACAACTTTATCGCCGCCATC,

R’ AGCCCTCCAGTCATTCACAA),

ENSDARG00000099980/bpifcl (F’ CAGAAGCAGATGAAGTTCATTAGTTCATTA,

R’ CTCCATGTTAGTGACTGCTTGTTGG)

### Hybridization chain reaction (HCR) *in situ*

HCR probes were designed and purchased from molecular instruments, or designed using the in-situ-probe-generator_v.0.3.2^22^ and purchased from IDT. The molecular instruments protocol for fixed whole-mount zebrafish embryos and larvae was followed with a few modifications. Whole fish were fixed in 4% paraformaldehyde (PFA) for 2 hours at room temperature, with a cavity opened for better PFA penetration, then organs dissected and rinsed in 1x phosphate-buffered saline. The methanol steps were eliminated and organs directly underwent proteinase K treatment which ranged from 30-50µg/ml for 10-15 minutes depending on the organ. 4pmol of each probe was utilized for hybridization. Organs were incubated with 1μg/ml of DAPI in 5x SSCT for one hour before a final 5x SSCT only wash step.

### Generation of constructs and transgenic lines

The RiboTag and AngioTag constructs were generated using Gateway Technology^23^. Rpl10a was PCR amplified from 24hpf zebrafish cDNA using forward primer GTG AGA GGG GAG ATA TCA CG and reverse primer CTA AGC GTA ATC TCC AAC ATC GTA TGG GTA GTA GAG GCG CTG TGG TTT TCC CAT G, and TOPO TA cloned into the PCR-II vector. Bridging PCR was then used to add EGFP and viral 2A sequence^24^ to generate the “RiboTag” construct pME-egfp-2a-rpl10a-3xHA. Gateway LR reactions were used to combine pDEST-IsceI-Flk7kb, pME-egfp-2a-rpl10a-3xHA, and p3E-polyA. The kdrl:egfp-2a-rpl10a3xHA “AngioTag” construct was digested with IsceI enzyme and microinjected into the blastomere of one-cell stage zebrafish embryos. A stable *Tg(kdrl:egfp-2a-rpl10a3xHA)^530^*^530^ germline transgenic line was established by screening through multiple generations.

The UAS:RiboTag construct was generated using SLiCE technology^25^. The pT1ump-14xUAS-MCS-POUT^26^ was digested with EcoRI and XhoI and then forward primer TCC CAT CGC GTC TCA GCC TCA CTT TGA GCT CCT CCA CAC GAA TTC GCC ACC ATG GTG TCA AAA G and reverse primer ACA TGT TCA GTT AAC GGT GGC TGA GAC TTA ATT ACT AGT CTC GAG TTA AGC GTA ATC TGG AAC ATC were used to slice clone the RiboTag cassette downstream from the 14x UAS sequence, using pME-egfp-2a-rpl10a-3xHA as the template.

The Gene C (ENSDARG00000098293) and Gene B (ENSDARG00000076721) mutants were generated using CRISPR-Cas9 technology, as outlined in Gagnon JA, et al^27^. For ENSDARG00000098293, the guide primer sequence was TAATACGACTCACTATAGGAATTGGGCGACTTACTGCGTTTTAGAGCTAGAAATAGC AAG, for ENSDARG00000076721, the guide primer sequence was TAATACGACTCACTATAGGTTTGGACCTCATGAGAGTGTTTTAGAGCTAGAA.

Genotype was determined using an ABI 3130 (Applied Biosystems) with screening primers TGTAAAACGACGGCCAGTATGGCTGTAGATGAATGAAGACT and GTGTCTTTCTCAGCACATGGTCAGAGG for ENSDARG00000098293 and screening primers TGTAAAACGACGGCCAGTCAGGTGTGTTTGGTGCTGAT and GTGTCTTCACGGGCATTAACTCACCAT for ENSDARG00000076721.

The Gene C mutant described in this manuscript has a 20bp deletion in exon 2.

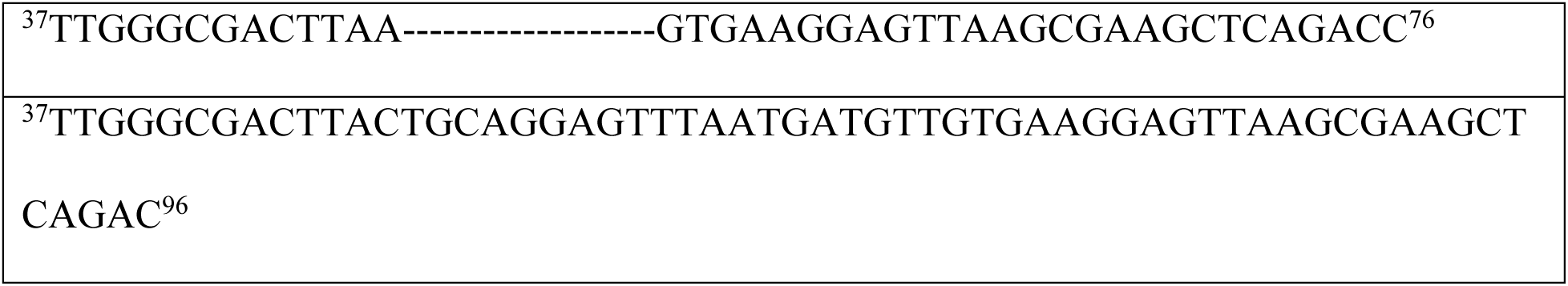

The Gene B mutant described in this manuscript has a 5bp deletion in exon 2.

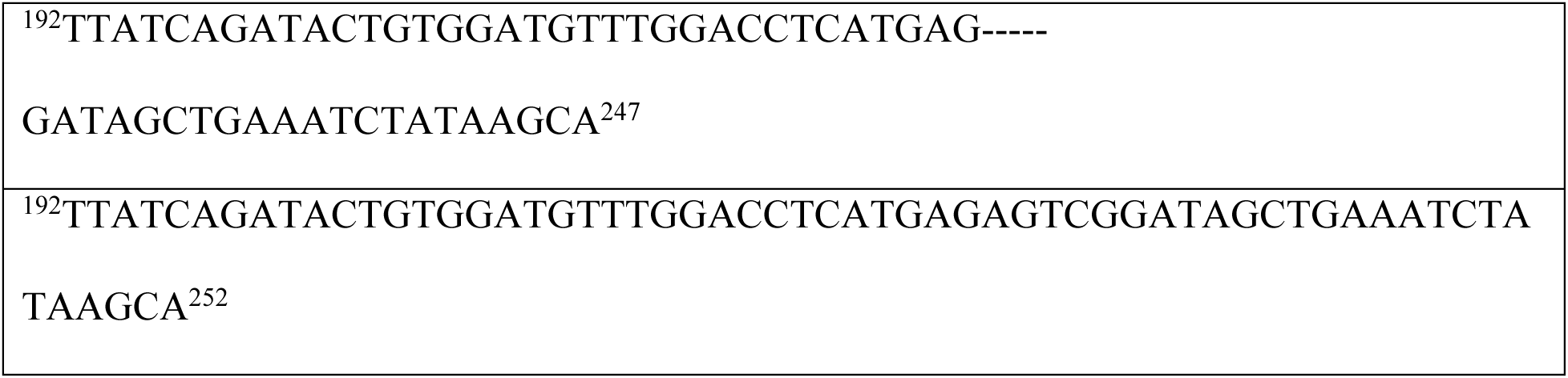

### Microscopic Imaging

Larvae used for imaging were anesthetized using 168mg/l Tricaine (1X Tricaine) and mounted in 0.8-1.5% low melting point agarose dissolved in embryo buffer and mounted on a depression slide. Confocal fluorescence imaging for larvae and adult organs was performed with an LSM 880 (Zeiss) or a Nikon TI2 with CSUW1 spinning disk microscope.

### Caudal Plexus Measurements

The dorsal-ventral width of the caudal plexus was measured under the first ISV posterior to the anal pore, using Fiji^28^. Measurements to the left, underneath, and to the right of the ISV were taken, and then averaged and normalized to the overall dorsal-ventral height of the embryo at the first ISV posterior to the anal pore, which was measured in a similar manner to the caudal plexus. The values for each genotype were then averaged, normalized to the wild type measurement, and the standard error of the mean was determined. A student’s t-test was run to determine significance.

### Study Approval

Zebrafish husbandry and research protocols were reviewed and approved by the NICHD Animal Care and Use Committee at the National Institutes of Health. All animal studies were carried out according to NIH-approved protocols, in compliance with the *Guide for the Care and use of Laboratory Animals*.

## RESULTS

### Translating ribosome affinity purification (TRAP) in zebrafish embryos

To determine whether Translating Ribosome Affinity Purification (TRAP) could be used to isolate polysome-bound mRNAs from developing zebrafish, we designed an egfp-2a-rpl10a3xHA RiboTag cassette. This cassette includes 60S ribosomal protein l10a (rpl10a) fused to triplicate hemagglutinin (HA) epitope sequences (rpl10a3xHA). To visualize transgene expression, the cassette also includes EGFP linked to rpl10a3xHA via a viral 2A cleavage site^24^, permitting bicistronic expression of separate EGFP and rpl10a3xHA polypeptides (**Figure 1A**). Assembly of Rpl10a3xHA protein into functional ribosomes marks translating mRNAs in cells expressing this transgene (**Figure 1B**). To test whether the transgene could be employed to isolate tagged polysomes using TRAP, we injected RiboTag mRNA into single-cell embryos, raised the embryos to 24hpf, homogenized them, and affinity-purified HA-tagged polysomes using αHA antibody followed by direct immunoprecipitation with Dynabeads® (**Figure 1C**). Western blotting of starting lysates, post-immunoprecipitation supernatants, and eluates from αHA pull-downs (**Figure 1D**) showed that Rpl10a3xHA protein was readily detected in RiboTag-injected (lane 2) but not in control uninjected (lane 1) embryos, and that Rpl10a3xHA protein was efficiently pulled down using the αHA antibody (lane 6) and depleted from the supernatant (lane 5).

**Figure 1.**
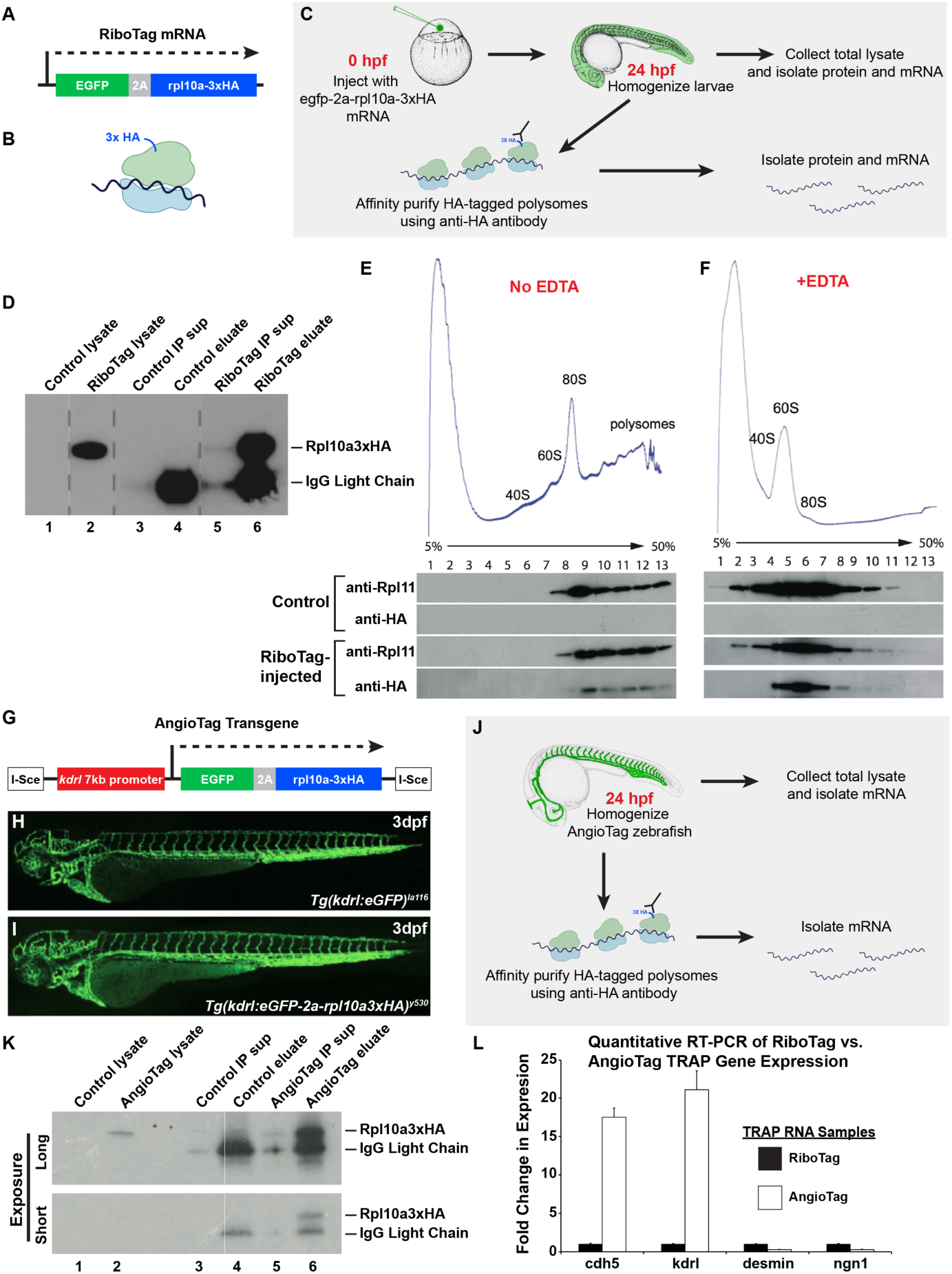
Affinity-tagged ribosomal protein subunits (Ribotags) can be used to purify translated RNAs *in vivo*. **(A)** Schematic diagram of the egfp-2a-rpl10a3xHA RiboTag cassette. **(B)** Schematic diagram of a 3xHA-tagged ribosome translating an mRNA. **(C)** Schematic diagram illustrating the workflow for TRAP purification of RNAs from RiboTag mRNA-injected zebrafish. **(D)** Western blot of starting lysates and TRAP supernatants and eluates from either control or RiboTag-injected animals, probed with αHA antibody. **(E)** Western blots of fractions collected from sucrose density gradient sedimentation of lysates from control or RiboTag-injected animals, probed with either αRpl11 or αHA. HA-tagged ribosomes sediment together with Rpl11-positive polysomes in the RiboTag-injected samples. **(F)** Western blots of fractions collected from sucrose gradient sedimentation of EDTA-treated lysates from control or RiboTag-injected animals, probed with either αRpl11 or αHA. Rpl11-positive endogenous ribosomes and HA-tagged ribosomes both sediment in the 60S fraction after EDTA treatment. **(G)** Schematic diagram of the IsceI(kdrl:egfp-2a-rpl10a3xHA) transgene. **(H)** Composite confocal micrograph of a 3dpf *Tg(kdrl:egfp)^la^*^116^ transgenic animal, for comparison purposes. **(I)** Composite confocal micrograph of a 3dpf *Tg(kdrl:egfp-2a-rpl10a3xHA)^530^*^530^ (“AngioTag”) transgenic animal. **(J)** Schematic diagram illustrating the workflow for TRAP purification of RNAs from AngioTag zebrafish. **(K)** Western blot of starting lysates and TRAP supernatants and eluates from either control or AngioTag animals, probed with αHA antibody, showing longer (top) and shorter (bottom) exposures of the same blot. IgG light chain is present in the pull-downs. **(L)** Quantitative RT-PCR measurement of the relative expression of *cdh5, kdrl, desmin,* and *ngn1* in cDNA samples prepared from TRAP purified RNA from either RiboTag control (black columns) or AngioTag (white columns) animals, showing enrichment of vascular specific genes and depletion of non-vascular specific genes in the AngioTag TRAP samples.

To ensure that Rpl10a3xHA protein is properly incorporated into the 60S ribosomal subunit, the 80S ribosome, and polysomes, we prepared lysates from RiboTag mRNA-injected and control-uninjected embryos and performed velocity sedimentation in sucrose density gradients, followed by Western blotting of collected fractions (**Figure 1E-F**). We probed the blots using an antibody recognizing endogenous Rpl11, a component of the 60S subunit, and found that Rpl11 sediments similarly within the 60S, 80S, and polysome fractions in either uninjected controls or RiboTag-injected animals (**Figure 1E**). When we probed the blots using an αHA antibody, we observed similar sedimentation patterns for Rpl10a3xHA protein in the RiboTag-injected animals, leading us to conclude that the epitope tag does not interfere with assembly of the 60S subunit, 80S ribosome, or polysomes (**Figure 1E**). As expected, disassembly of polysomes using EDTA treatment causes both Rpl11 and Rpl10a3xHA to elute only in the 60S fraction; they are not seen in the 80S or polysome fractions (**Figure 1F**). These results show that Rpl10a3xHA protein assembles into ribosomes *in vivo* and can be used for isolation of tagged polysomes from zebrafish.

### Isolating endothelial-specific RNA using AngioTag zebrafish

We next set out to create transgenic zebrafish expressing the RiboTag construct specifically within endothelial cells. We utilized the previously published endothelial specific 7kb *kdrl* promoter^16^ to design a IsceI(kdrl:egfp-2a-rpl10a3xHA) AngioTag transgene expressing the RiboTag cassette in an endothelial cell type-specific manner (**Figure 1G**). We obtained a strongly expressing *Tg(kdrl:egfp-2a-rpl10a3xHA)^530^*^530^ germline transgenic line that shows a vascular-restricted EGFP expression closely matching the expression pattern of a previousy generated *Tg(kdrl:egfp)^la^*^116^ ^16^ transgenic line driving EGFP alone (**Figure 1H-I**). To test whether this AngioTag transgenic line could be used for TRAP enrichment of endothelial mRNAs, we prepared homogenates from 24hpf AngioTag or control nontransgenic animals and subjected the lysates to the TRAP procedure (**Figure 1J**). Western blotting of starting lysates (lanes 1 and 2), TRAP supernatants (lanes 3 and 5), and TRAP eluates (purified tagged polysomes, lanes 4 and 6) from non-transgenic (lanes 1,3,4) or AngioTag transgenic (lanes 2,5,6) animals (**Figure 1K**) showed that Rpl10a3xHA protein was readily detected in AngioTag (lane 2) but not in control (lane 1) non-transgenic total lysates, and that Rpl10a3xHA was pulled down from AngioTag lysates using αHA antibody (lane 6). To assess whether the AngioTag TRAP procedure resulted in selective enrichment of endothelial-specific mRNAs, we performed quantitative RT-PCR for endothelial and non-endothelial genes using RNA collected from either 24hpf AngioTag transgenic animals or 24hpf RiboTag mRNA injected animals. RNA isolated from AngioTag animals was highly enriched for endothelial-specific genes *(cdh5, kdrl)* and depleted of genes expressed specifically in non-vascular tissues *(desmin, ngn1)* compared to the RiboTag mRNA injected control RNA samples (**Figure 1L**).

### Expression profiling using TRAP-RNAseq in zebrafish

Having shown that our AngioTag transgenic line could be used to selectively enrich for endothelial gene expression, we wanted to compare AngioTag TRAP-RNAseq endothelial expression profiling to FACS endothelial expression profiling using previously published methods in zebrafish^29–31^ (**Figure 2A-B**). As noted above, FACS protocols require the dissociation of an embryo into a single cell suspension, disrupting cell-cell, cell-matrix, and other external interactions for a prolonged period of time prior to cell lysis and RNA collection (**Figure 2A**). In contrast, the TRAP procedure involves immediate homogenization, cell lysis, and stabilization of translating RNAs (**Figure 2B**), providing an instant ‘snapshot’ of translated gene expression at the time of embryo homogenization, one that is potentially more representative of an ‘*in vivo’* profile.

**Figure 2.**
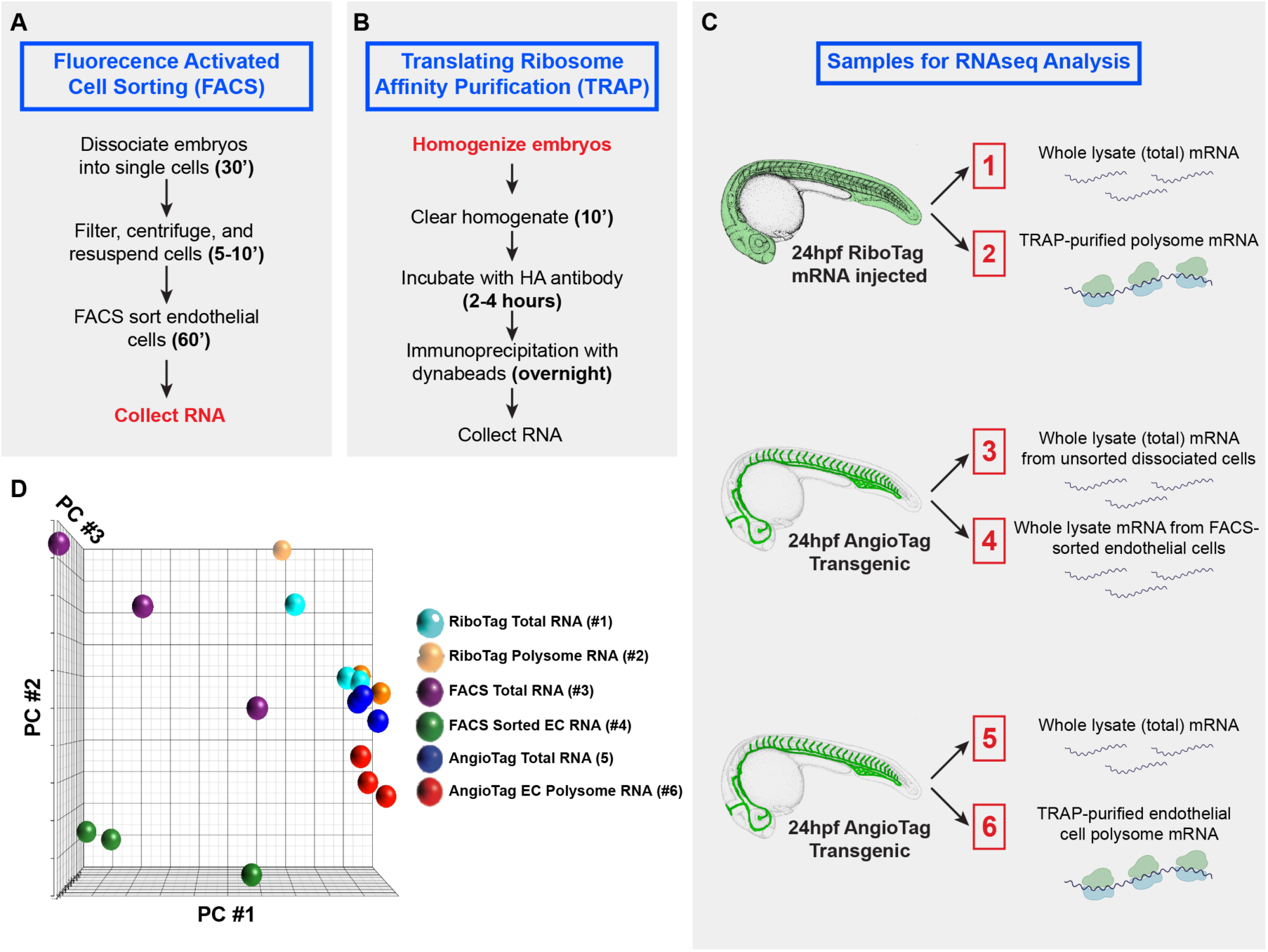
Comparative workflow for preparation of endothelial-specific RNAs using fluorescence activated cell sorting (FACS) versus translating ribosome affinity purification (TRAP) **(A)** Workflow for FACS of EGFP-positive endothelial cells and RNA preparation from 24hpf *Tg(kdrl:egfp-2a-rpl10a3xHA)^530^*^530^ AngioTag transgenic animals (sample #4 in panel C). RNA is collected from the cells after more than 1½ hours of embryonic dissociation and cell sorting. **(B)** Workflow for TRAP of translated mRNAs from 24hpf egfp-2a-rpl10a3xHA RiboTag mRNA-injected animals (sample #2 in panel C) or *Tg(kdrl:egfp-2a-rpl10a3xHA)^530^*^530^ (AngioTag) transgenic animals (sample #6 in panel C). Lysate are prepared from intact animals and the RNA is stabilized immediately at the beginning of the procedure. **(C)** Samples collected in triplicate for RNAseq analysis. 24hpf RiboTag mRNA injected (samples 1 and 2), dissociated AngioTag transgenic (samples 3, 4), or AngioTag transgenic animals (samples, 5, 6). Part of each sample was used for whole lysate total RNA collection (samples 1, 3, and 5). The remainder of each sample was used for either TRAP purification of total (sample 2) or endothelial (sample 6) polysome mRNA, or for FACS sorting of EGFP-positive endothelial cells followed by RNA isolation (sample 4). **(D)** Principal component analysis of RNAseq data obtained from the six sample types noted in panel C, each run in triplicate (total of 18 samples).

We performed RNAseq in triplicate on a total of six different sample types (**Figure 2C**). RNA was prepared from TRAP-purified polysomes from 24hpf RiboTag mRNA injected animals (Sample 2, ‘whole TRAP translatome’), from FACS-sorted endothelial cells from 24hpf AngioTag transgenic animals (Sample 4, ‘endothelial FACS transcriptome’), and from TRAP-purified polysomes from 24hpf AngioTag transgenic animals (Sample 6, ‘endothelial TRAP translatome’). For each of these samples, we also collected control RNA from either the starting lysates used for TRAP (Samples 1 and 5) or from unsorted cells (Sample 3). Three experimental replicates were performed for each of these sample types from separately collected samples, for a total of 18 samples used for RNAseq analysis. A minimum of 45 million reads were obtained for each sample, with an average of 84% of reads mapping uniquely (**Supplemental Table 1**). All sequence data was deposited with GEO (accession # GSE121973). Principal component analysis (**Figure 2D**) showed that the RiboTag-TRAP (1,2) and AngioTag-TRAP (5,6) starting lysates and TRAP samples clustered fairly closely together, while the FACS unsorted (3) and sorted (4) cell samples showed considerable divergence both from the other samples and from one another.

Comparing the whole-embryo lysate transcriptome (Sample 1) and matched whole-embryo TRAP translatome (Sample 2) from animals ubiquitously expressing RiboTag (**Figure 3A**) showed that 1460 genes were significantly differentially represented in the two RNAseq data sets (Benjamini-Hochberg (BH)-adjusted p < 0.05), with 829 genes showing increased representation (log2(fold)>0.5) and 631 genes showing reduced representation (log2(fold)<-0.5) in the whole-embryo TRAP translatome (Sample 2) vs. transcriptome (Sample 1) datasets. Nine of the ten PANTHER overrepresentation test^32^ GO terms with the largest fold increase in the whole animal translatome (Sample 2) vs. the transcriptome (Sample 1) were related to splicing and/or RNA metabolism, suggesting mRNAs for these genes may be more highly translated (**Figure 3B**). To determine whether there are any sequence features associated with increased or decreased translation, we examined relative codon usage associated with genes over- or under-represented in sample 2 vs. sample 1, identifying strong preferences for and against particular codons in more-versus less-highly translated genes (**Figure 3C**). Interestingly, our findings closely align with previously published data examining codon usage bias in highly transcribed genes in the zebrafish genome^33^. We also examined the 3’UTR sequence length and found that more highly translated genes have, on average, shorter 3’UTRs (**Figure 3D**), in agreement with prior work suggesting mRNAs with long 3’UTRs have increased numbers of miRNA binding sites leading to lower levels of translation^34,35^.

**Figure 3.**
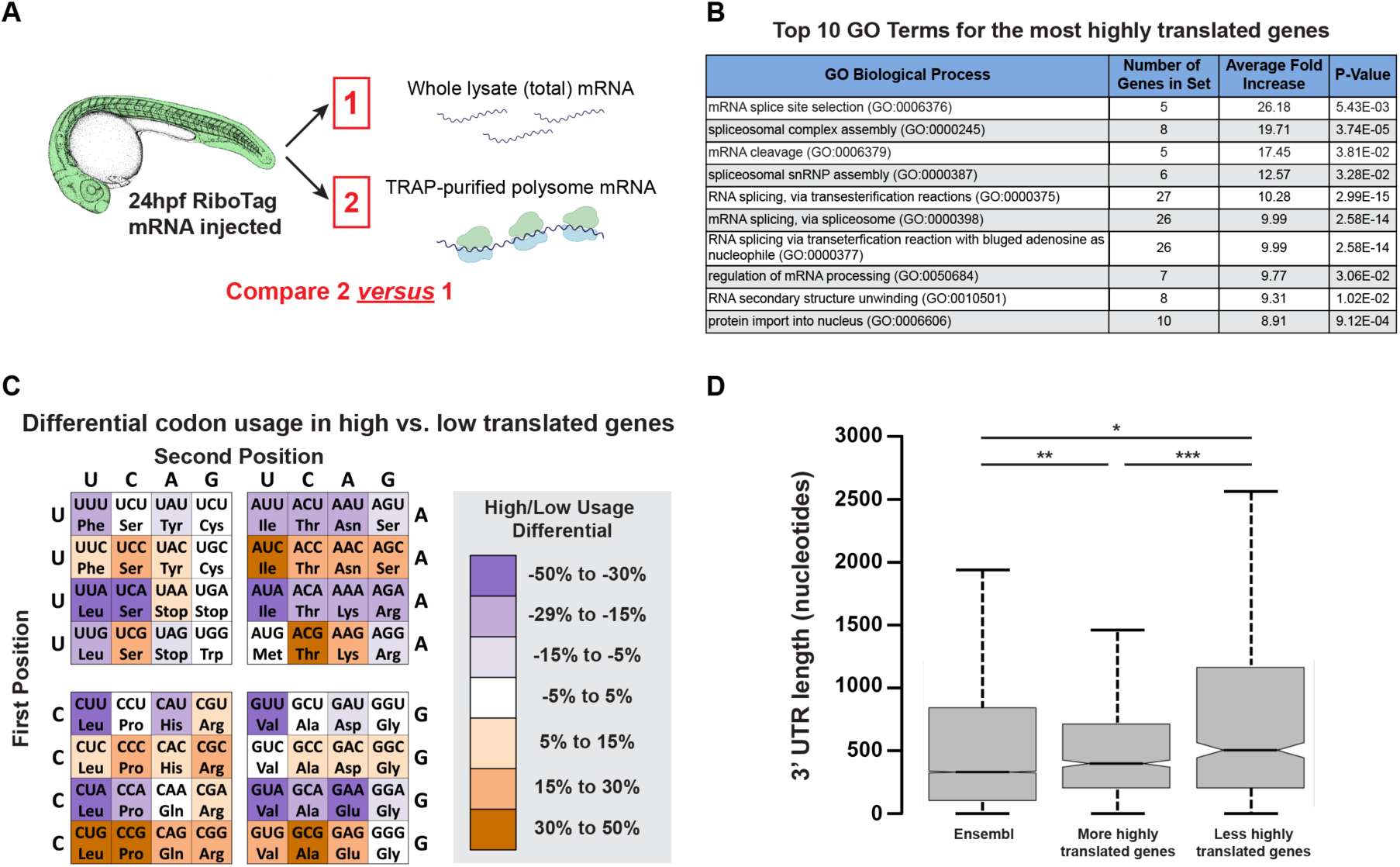
Comparison of the whole-animal transcriptome to its translatome. **(A)** Samples collected for RNAseq analysis of whole animal translatome (sample 2) vs. whole animal transcriptome (sample 1) from 24 hpf RiboTag mRNA injected animals. **(B)** Top 10 GO terms for the highest-translated genes based on average fold increase. **(C)** Percentage fold-change in the percentage usage of each codon between the most highly translated genes (fold ≥ 1.5, BH-adjusted p ≤ 0.05 comparing samples 2 vs. 1) and the least highly translated genes (fold ≤ 1.5, BH-adjusted p ≤ 0.05 comparing samples 2 vs. 1). The color-coding scheme shows the percentage change in the relative usage of each codon between the two sets of genes (with the usage of each codon calculated as a percentage of the total number of codons coding for a particular amino acid). **(D)** 3’ UTR length of all Ensembl genes (first column) compared to the most highly translated (second column) and least highly translated (third column) sets of genes.

### AngioTag profiling endothelial genes

Having shown that TRAP-RNAseq can be used effectively to profile translating mRNAs *in vivo* in the zebrafish, we next sought to examine whether our AngioTag transgenic line could be used for specific profiling of endothelium *in vivo.* As noted above, FACS has been used previously to isolate endothelial cells from developing zebrafish for gene expression profiling^29–31^ so, for purposes of comparison, we began by carrying out RNAseq analyses on mRNA from endothelial cells isolated from 24hpf AngioTag animals by FACS (**Figure 4A-C**). RNAseq experiments were carried out on mRNA isolated from total dissociated cells from 24hpf AngioTag embryos (Sample 3) and on mRNA from EGFP-positive endothelial cells FACS-sorted from the same dissociated cell population (Sample 4; **Figure 4A**). Comparative analysis of genes represented in the two RNAseq data sets showed that 3360 genes were significantly enriched in the FACS-sorted endothelial cells (log2(fold)>0.5, BH-adjusted p ≤ 0.05), while 3174 genes showed significant depletion in the FACS-sorted endothelial cell population compared to the total dissociated cell population (log2(fold)<-0.5, BH-adjusted p ≤ 0.05; **Figure 4B**). Six out of the top twenty PANTHER overrepresentation test GO terms showing the highest fold enrichment in FACS-sorted endothelial cells were endothelial/vascular-related terms, while the other 14 most highly enriched GO terms were non-endothelial (**Figure 4C**).

**Figure 4.**
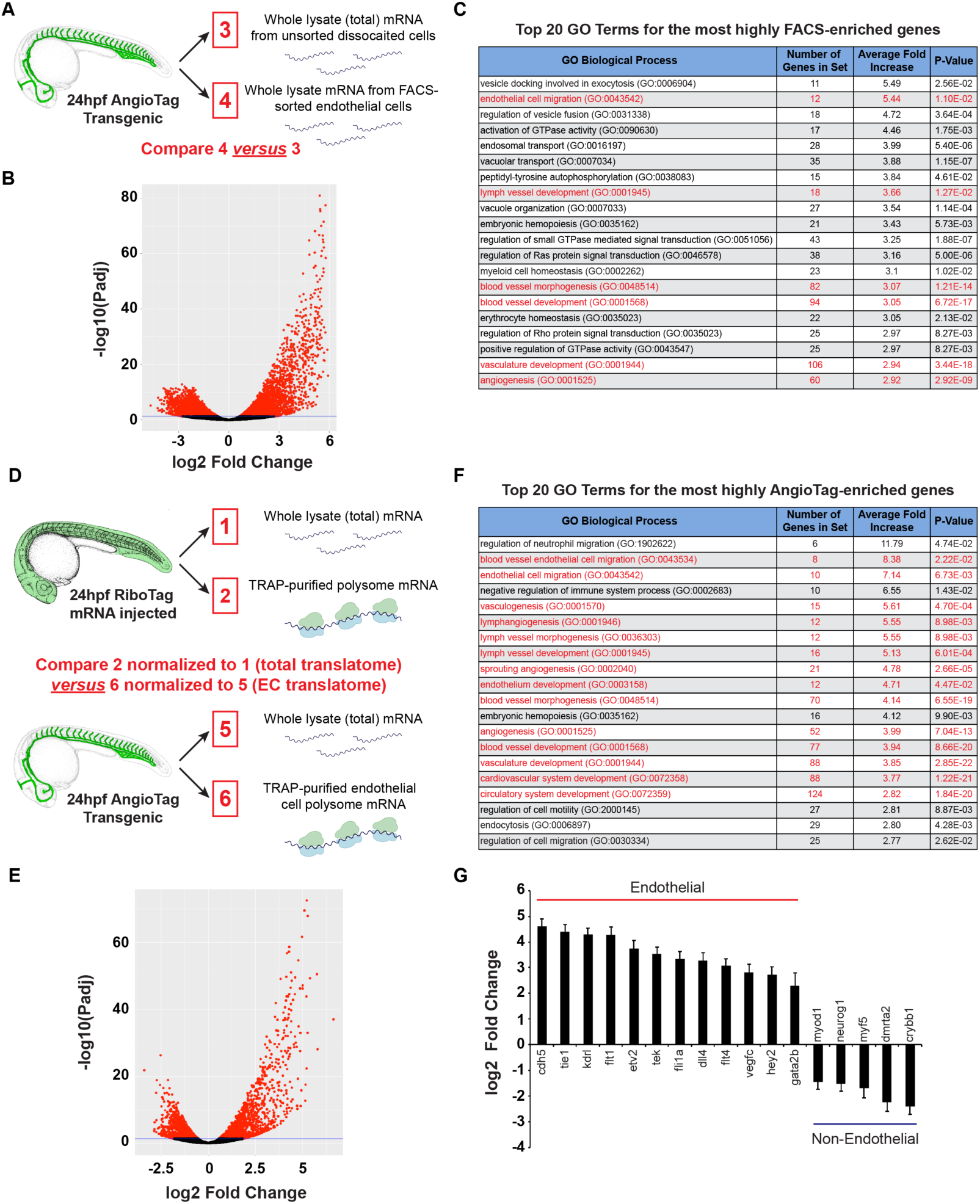
Translating Ribosome Affinity Purification shows greater endothelial gene enrichment compared to Fluorescence Activated Cell Sorting. **(A)** Samples collected from 24 hpf AngioTag transgenic animals dissociated into a cell suspension, for RNAseq analysis of mRNA from either FACS sorted endothelial cells (sample 4) or unsorted input cells (sample 3). **(B)** Volcano plot of samples 4 vs. 3. **(C)** Top twenty GO terms for genes most highly enriched in the FACS sorted endothelial cell samples vs. whole animal unsorted cells. Six out of the top twenty GO terms represent endothelial process-related terms (in red text). **(D)** Samples collected for RNAseq analysis of TRAP purified endothelial cell polysome mRNA from 24 hpf AngioTag transgenic animals (sample 6) normalized to its input total mRNA (sample 5) compared to TRAP purified total embryonic polysome mRNA from 24 hpf RiboTag mRNA injected animals (sample 2) normalized to its input total mRNA (sample 1). **(E)** Volcano plot of endothelial enrichment using the sample comparison shown in panel D. **(F)** Top twenty GO terms for genes most highly enriched in the TRAP purified endothelial polysome mRNA samples. Fourteen out of the top twenty GO terms represent endothelial process-related terms (in red text) compared to the only six seen with FACS sorted endothelial cells in panel C. **(G)** Log2 fold enrichment of endothelial specific genes and depletion of non-endothelial specific genes in AngioTag TRAP-RNAseq sample 6 (normalized to 5) as compared to RiboTag TRAP-RNAseq sample 2 (normalized to 1).

Next, we examined the results of profiling the endothelium in the AngioTag line using TRAP-RNAseq (**Figure 4D-G**). We compared RNAseq data sets of whole animal translatome (Sample 2) and endothelial translatome (Sample 6) after normalization of each of these samples to their whole-animal starting lysate transcriptome (samples 1 and 5, respectively; **Figure 4D**). 2136 genes were enriched in the endothelial translatome (log2(fold)> 0.5, BH-adjusted p ≤ 0.05), while 1867 genes show some depletion when compared to the whole animal translatome (log2(fold) ≤ - 0.5, BH-adjusted p ≤ 0.05; **Figure 4E**). Of the top twenty GO terms enriched in the endothelial TRAP-RNAseq dataset, fourteen were endothelial/vascular-related terms (**Figure 4F**), a substantially higher number than found in the FACS-sorted endothelial cell data (**Figure 4C**). Although six of these fourteen GO terms were also present in the top twenty terms for the FACS-sorted endothelial cells, they showed greater fold-enrichment with lower p-values in the TRAP-RNAseq dataset (**Supplemental Table 2**). As expected, several known endothelial-specific genes are highly enriched in the AngioTag TRAP-RNAseq data set including *kdrl*, *cdh5*, *tie1*, and *flt1*, while genes that are not expressed in endothelium are depleted (**Figure 4G**).

The vast majority of genes highly enriched by AngioTag TRAP-RNAseq are annotated, and many of these genes are already known to be expressed in the endothelium. However, our enriched gene set included genes not previously reported to be expressed in endothelium (**Figure 5A-B**). *In situ* hybridization for four of these genes, *scpp8, exoc3l2a, slc22a7b.1,* and *bpifcl,* confirmed their endothelial-specific expression pattern (**Figure 5C-F**), suggesting that these genes may be involved in vascular development. In addition to annotated genes, our enriched gene set also includes unannotated genes lacking a previously reported link to endothelium, including four genes found amongst the 50 most highly enriched genes in our dataset (**Figure 5G**), each of which shows at least 20-fold endothelial enrichment (**Figure 5H**). Whole mount *in situ* hybridization confirms that all four of these genes have highly endothelial-specific expression patterns (**Figure 5I-P**), with expression of one of the genes largely restricted to the endothelium of the caudal vascular plexus (**Figure 5K-L**). To explore the functional significance of a few of the unannotated genes, we use CRISPR/Cas9 technology to generate targeted mutations in Gene B (ENSDARG00000076721) and in Gene C (ENSDARG00000098293). We isolated a 5 base pair deletion mutant in Gene B (76721^y587^) generating a polypeptide truncated after 77 amino acids (**Figure 6A**), and a 20 base pair deletion mutant in Gene C (98293^y588^) coding for a polypeptide truncated after 25 amino acids (**Figure 6F**). Homozygous 76721^y587^ mutants display a mild enlargement of the caudal vascular plexus (**Figure 6B-E**), while homozygous 98293^y588^ mutants show reduced growth of mid- and hindbrain cranial central arteries (**Figure 6G-I**). These results suggest that at least two of the four novel, unannotated genes from this study have roles in early vascular development.

**Figure 5.**
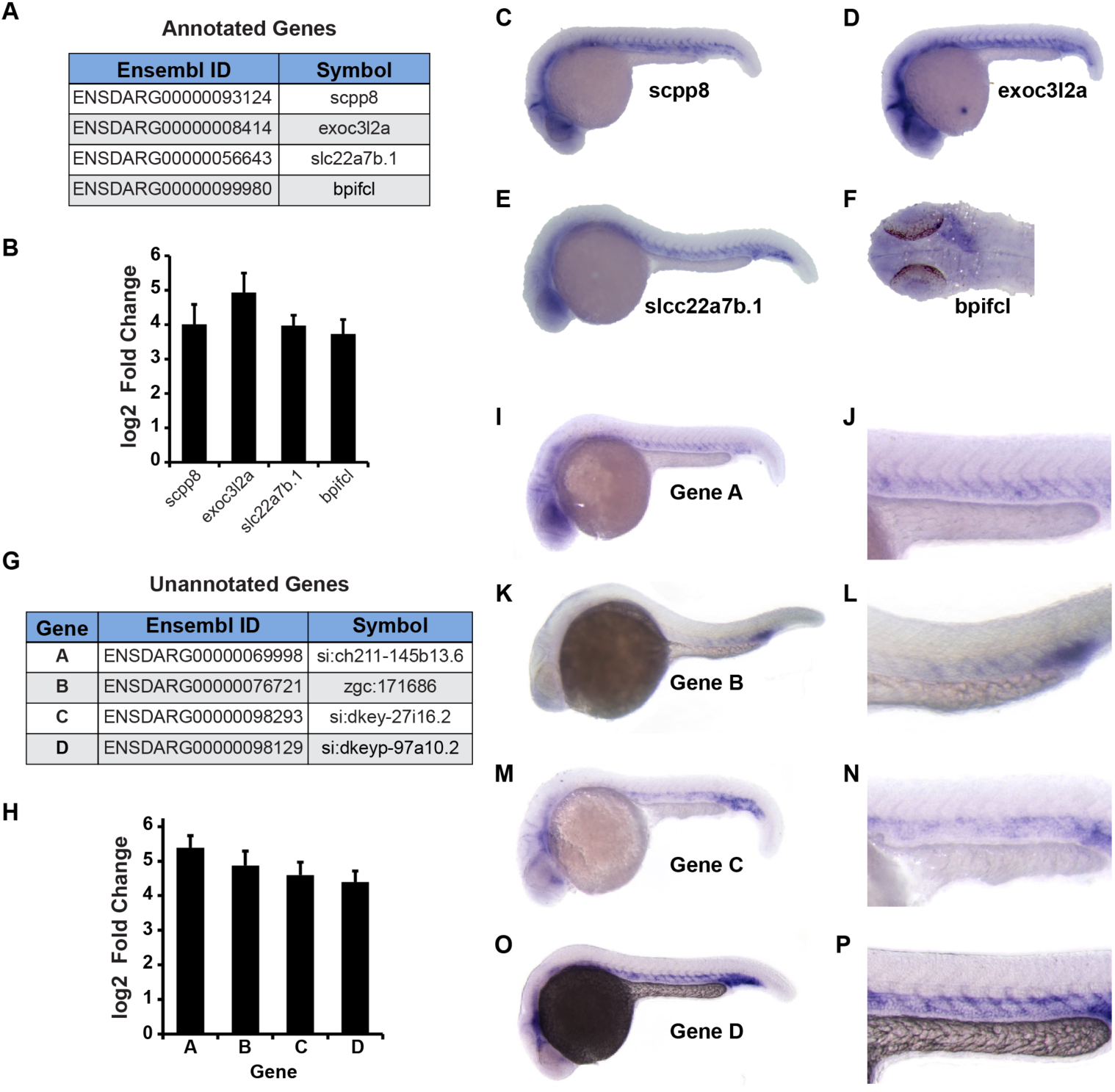
TRAP-RNAseq reveals novel endothelial genes expressed in early vascular development. **(A)** Four annotated genes found among the top 50 genes enriched in the AngioTag TRAP-RNAseq data set (sample 6) without previously recognized endothelial expression. **(B)** Log2 fold enrichment of the four annotated genes noted in panel A in AngioTag TRAP-RNAseq sample 6 (normalized to 5) compared to RiboTag TRAP-RNAseq sample 2 (normalized to 1). **(C-F)** Whole mount *in situ* hybridization of 24 hpf wild type zebrafish probed for *scpp8* (C), *exoc312a* (D), *slcc22a7b.1* (E), and *bpifcl* (F). Images shown are either lateral views of the whole animal (panels C-E), or a dorsal view of the head with staining of the heart noted (F). **(G)** Four unannotated genes found among the top 50 genes enriched in the AngioTag TRAP-RNAseq data set (sample 6). **(H)** Log2 fold enrichment of the four unannotated genes noted in panel G in AngioTag TRAP-RNAseq sample 6 (normalized to 5) compared to RiboTag TRAP-RNAseq sample 2 (normalized to 1). **(I-P)** Whole mount *in situ* hybridization of 24 hpf wild type zebrafish probed for unannotated gene A (panels I-J), gene B (panels K-L), gene C (panels M-N), and gene D (panels O-P). Images shown are lateral views of the whole animal (panels I,K,M,O), with higher magnification lateral views of the trunk (panels J,L,N,P).

**Figure 6.**
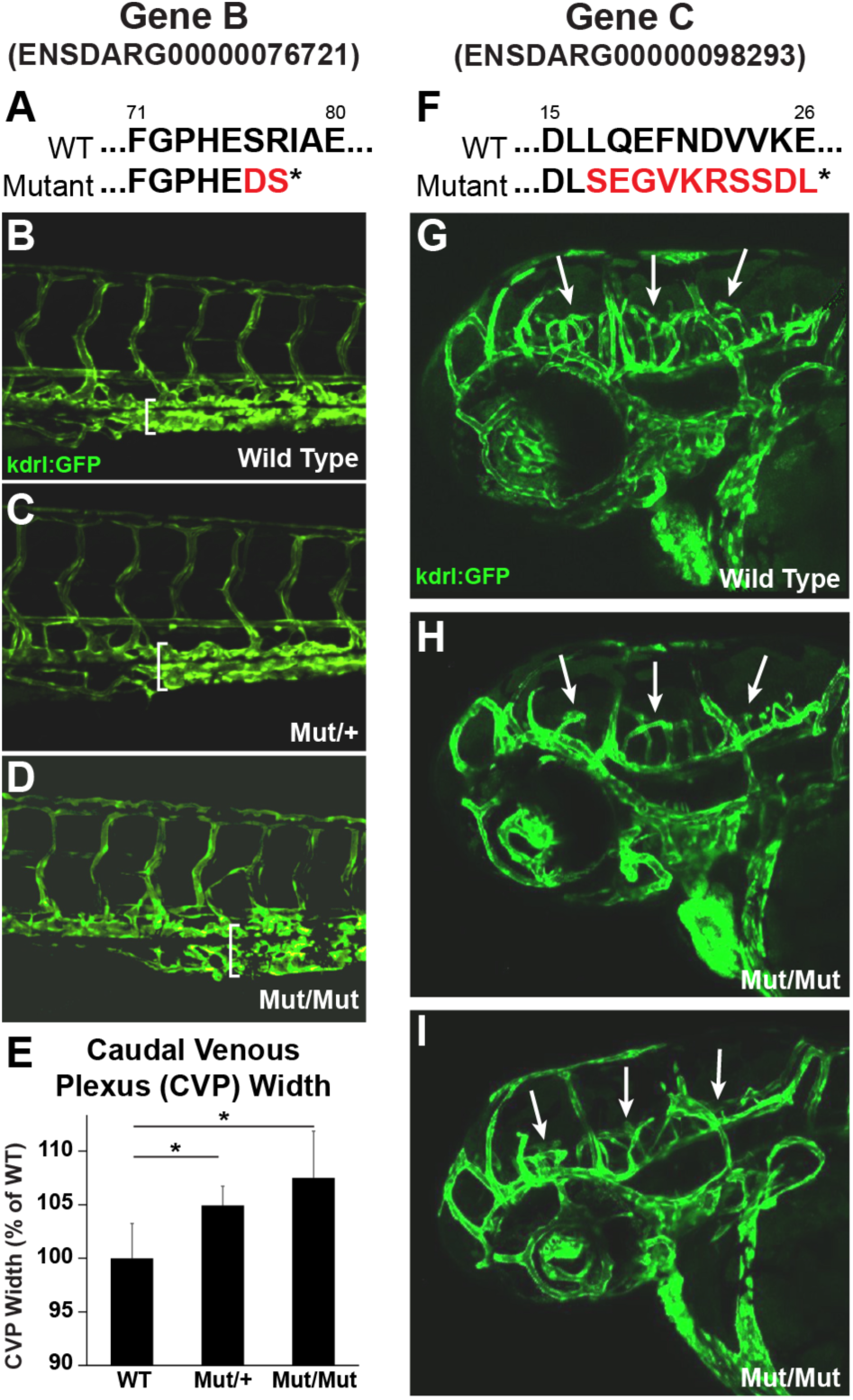
Mutants of unannotated endothelial genes have vascular phenotypes. **(A)** In *76721^530^*^587^ mutants a 5 bp deletion in ENSDARG00000076721 introduces an early stop codon, resulting in a protein truncation from 316 amino acids to 78 amino acids. **(B-D)** Confocal microscopy of 3dpf embryos from a *76721^530^*^587^*^/+^* heterozygous incross showing a dilated caudal plexus in heterozygous (C) and homozygous mutant (D) embryos as compared to wild type siblings (B). **(E)** Graph comparing the caudal plexus height of wild type, *76721^530^*^587^*^/+^* heterozygous, and *76721^530^*^587^*^/y^*^587^ homozygous mutant siblings. Values shown are the percent change compared to wild type (N = 41 wild types, 58 heterozygotes, 23 homozygous mutants). **(F)** In *98293^530^*^588^ mutants a 20 bp deletion in ENSDARG00000098293 introduces an early stop codon, resulting in a protein truncation from 104 amino acids to 27 amino acids. **(G-I)** Confocal microscopy of 3dpf wild type (G) and *98293^530^*^588^*^/y^*^588^ homozygous mutant embryos (H-I) reveals decreased cranial vasculature in mutant embryos compared to wild type (white arrows). * student’s t-test p-value ≤ 0.05.

### AngioTag profiling of adult zebrafish organs

In addition to profiling genes involved in vascular development we wanted to know if our AngioTag transgenic fish could be used to reveal tissue-specific differences in the endothelial translatomes of different adult organs (**Figure 7**). To test this we carried out TRAP-RNAseq on skin, muscle, liver, heart, and brain dissected from adult AngioTag transgenic fish, and on a whole fish control. Each tissue sample was run in triplicate and each replicate contained organs from four adult fish. Angiogenesis and blood vessel morphogenesis were among the top ten GO terms for each organ’s TRAP pulldown compared to their whole organ total input (**Supplemental Tables 3-7**). To uncover endothelial genes common to all vascular beds, we compared the endothelial translatome of each organ (IP) to its whole organ total (input) mRNA, then looked for genes that showed significant enrichment across all of the organ datasets (log2(fold) ≥1.2, padj<0.05) (**Figure 7A**). As expected we saw enrichment of well-known endothelial markers such as *kdrl*, *cdh5*, and several Notch pathway members (**Figure 7B**). We verified robust expression of *kdrl* and *cdh5* in all five organs using the *Tg(kdrl:egfp)^la^*^116^ transgenic line and whole mount hybridization chain reaction (HCR) *in situs* for *cdh5* (**Figure 7C-G**). Our AngioTag data also revealed strong cross-organ endothelial enrichment of some less well known vascular genes, including *fgd5a*, *bcar1,* and our unannotated Gene D (**Figure 7B**).

**Figure 7.**
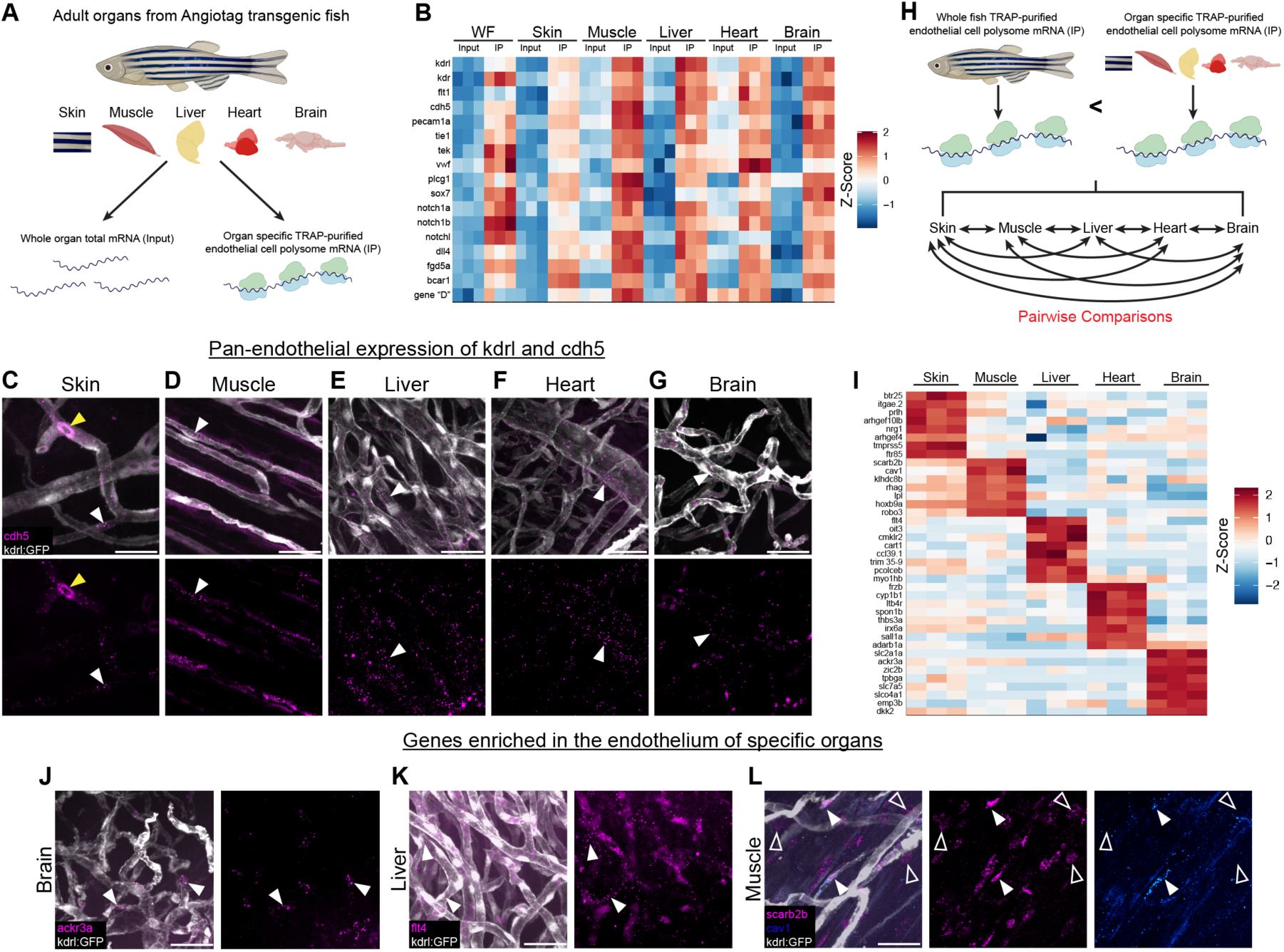
Endothelial profiling using TRAP-RNAseq reveals common vasculature signatures and unique gene expression profiles across vascular beds of different organs. **(A)** Samples collected for RNAseq analysis of TRAP purified endothelial cell polysome mRNA from five different organs, skin, muscle, liver, heart, brain, and whole fish of AngioTag transgenic animals normalized to each organ’s input total mRNA. **(B)** Heatmap displaying common endothelial genes enriched in the vasculature of the whole fish (WF) and across vascular beds of each organ compared to their input controls colored by their gene-wise z-scores of the log-transformed normalized counts. TRAP pulldown (IP) and total organ mRNA (input) for each replicate for each organ are shown. **(C-G)** Confocal images of adult *Tg(kdrl:egfp)^la^*^116^ transgenic fish with hybridization chain reaction (HCR) *in situ* of *cdh5* showing expression of *kdrl* (white) and *cdh5* (magenta) in the vasculature of the skin (C), muscle (D), liver (E), heart (F), and brain (G). Merged and *cdh5* only channels are shown. White arrowheads indicate *cdh5* transcript expression while yellow arrowheads show blood cell autofluorescence. **(H)** Schematic depicting screening steps used to determine genes unique to the vascular beds of each organ. To be included genes first had to be enriched in the vasculature of an organ compared to its total tissue and vascular expression in that organ had to be greater than vascular expression in the whole fish. Genesets then underwent pairwise comparisons between each organ to determine if they were shared or unique to a particular organ. **(I)** Heatmap displaying unique endothelial genes enriched in the vascular beds of each organ colored by their gene-wise z-scores of the log-transformed normalized counts. TRAP pulldowns (IP) for each replicate for each organ are shown. **(J-L)** Confocal images of adult *Tg(kdrl:egfp)^la^*^116^ transgenic fish (white) with hybridization chain reaction (HCR) *in situ* of *ackr3a* (magenta) in the brain (J), *flt4* (magenta) in the liver (K), and *scarb2b* (magenta) and *cav1* (blue) in the muscle (L). White solid arrowheads indicate probe expression in the vessel, and white open arrowheads indicate probe expression in the muscle fiber. Scale bars are 25μm.

We carried out an additional two-step comparison to uncover genes uniquely enriched in the endothelium of specific organs (**Figure 7H**; see methods for full details). We began by first selecting all genes that showed significantly greater enrichment in the endothelial translatomes of specific organs than in the whole fish endothelial translatome (**Figure 7H**, top). The selected genes were then subjected to further pairwise comparisons between each of the organs to find those more highly enriched in one particular organ compared to all of the others (**Figure 7H**, bottom). For a gene to be considered unique to the endothelium of a particular organ it had to meet each of the following criteria: 1) endothelial gene expression is at least 1.5 log2(fold) greater than in the starting lysate of that organ, 2) endothelial enrichment in the most-enriched organ is at least 2 log2(fold) greater than in the next-most-enriched organ, and 3) the difference in endothelial gene expression between most-enriched organ and next-most-enriched organ is at least 2 log2(fold) greater then their tissue lysate differences. Applying these very strigent criteria resulted in a small list of genes uniquely enriched in the vasculature of each organ with the top 7-8 genes that fit these criteria for each organ being displayed (**Figure 7I**). As expected, the well known blood-brain barrier gene *slc2a1a*, also known as *glut1*, was highly expressed in the brain vasculature only. The *atypical chemokine receptor 3a* (*ackr3a*) gene was also specifically enriched in the brain vasculature, as validated using HCR *in situs* (**Figure 7I,J**). Interestingly, the top GO terms for liver endothelium were lymphangiogenesis, lymph vessel morphogenesis, and lymph vessel development, despite the fact that we were profiling blood vessels, not lymphatic vessels (**Supplemental Table 5**). In keeping with this finding, we found that the lympho-venous marker *flt4* was selectively enriched in the vasculature of the liver compared to other organs (**Figure 7I**). HCR *in situs* for *flt4* in liver tissue confirmed its colocalization with *Tg(kdrl:egfp)^la^*^116^ transgene-positive blood vessels (**Figure 7K**). The expression of lymphatic endothelial genes in the liver blood vasculature may be indicative of similarities in vessel function, since lymphatic capillaries are highly permeable and liver vessels are thought to be one of the most permeable types of blood vessels^36,37^. As has been previously reported using single-cell RNAseq methods in mice^38^, we also found organ-specific genes whose expression was shared between both the endothelium and surrounding cells. In muscle, the *scavenger receptor class B, member 2b* (*scarb2b*) gene, predicted to encode a lysosomal membrane protein^39^, was expressed in both muscle endothelial cells and in muscle fibers (**Figure 7L**), despite expression of other genes such as *caveolin 1* (*cav1*) being restricted to only muscle vessels **(Figure 7L**).

Together, our results suggest that TRAP-RNAseq using our AngioTag transgenic line is highly effective for *in vivo* profiling of endothelial gene expression during development as well as in the vascular beds of adult organs.

### A RiboTag reporter line for profiling any cell or tissue of interest

Our success in carrying out endothelial profiling using AngioTag TRAP-RNAseq led us to ask whether we could generalize this method to profile other non-endothelial tissues. In order to do this, we generated a *Tg(uas:egfp-2a-rpl10a2xHA)^530^*^531^ UAS:RiboTag transgenic line to drive expression of the RiboTag cassette in any tissue or cell type of interest for which a Gal4 driver line is available (**Figure 8A**). The UAS:RiboTag line showed strong, tissue-specific expression after being crossed to a variety of different Gal4 driver lines, including a muscle-specific *Tg(xa210:gal4)^530^*^241^ line^17^ (**Figure 8B**), an endothelial-specific *Tg(fli1a:gal4ff)^ubs^*^4^ line^18^ (**Figure 8C)**, and a neural-specific *Tg(huC:gal4)* line^19^ (**Figure 8D**). TRAP isolation of mRNA from embryos obtained from each of these crosses revealed strong enrichment of tissue-specific genes in samples prepared from the appropriate lines (**Figure 8E-G**), specifically *kdrl* in the endothelial line and *snap25* in the neural line. The enrichment of the endothelial-specific *kdrl* gene in TRAP-purified mRNA from *Tg(uas:egfp-2a-rpl10a2xHA)^530^*^531^; *Tg(fli1a:gal4ff)^ubs^*^4^ double-transgenic animals was comparable to that found in TRAP-purified mRNA from our *Tg(kdrl:egfp-2a-rpl10a3xHA)^530^*^530^ AngioTag line (compare **Figure 1L** to **Figure 8F**), suggesting that the UAS:RiboTag line provides a similarly effective tool for tissue-specific gene expression profiling.

**Figure 8.**
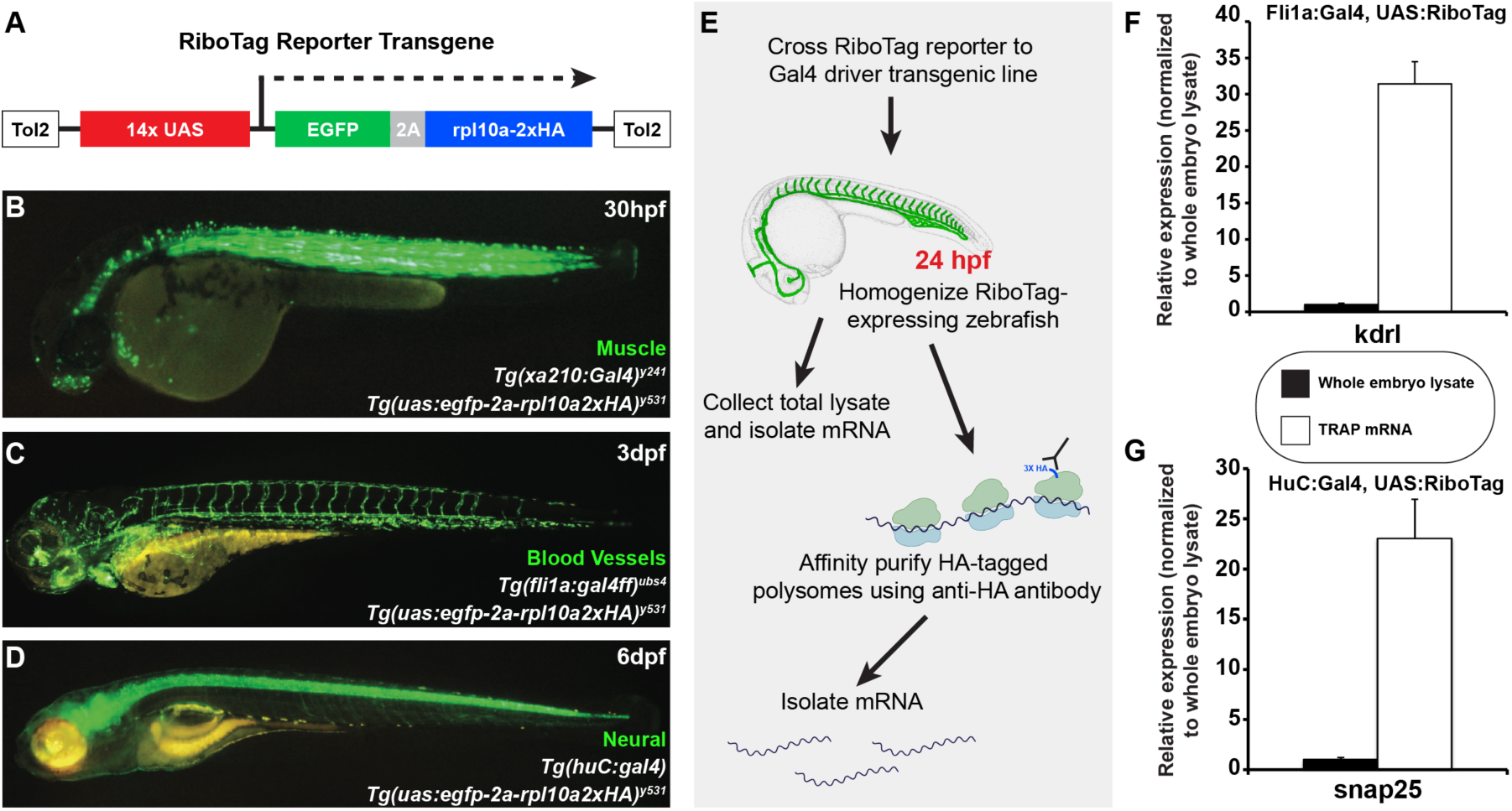
TRAP profiling can determine gene expression in different tissues and cell types using Ribotag Reporter transgenics. **(A)** Schematic diagram of the Tol2(uas:egfp-2a-rpl10a2xHA) RiboTag Reporter transgene. **(B)** Green epifluorescence photomicrograph of a 30hpf *Tg(uas:egfp-2a-rpl10a2xHA)^530^*^531^*; Tg(xa210:gal4)^530^*^241^ double transgenic animal. **(C)** Green epifluorescence photomicrograph of a 3dpf *Tg(uas:egfp-2a-rpl10a2xHA)^530^*^531^*; Tg(fli1a:gal4ff)^ubs^*^4^ double transgenic animal. **(D)** Green epifluorescence photomicrograph of a 6dpf *Tg(uas:egfp-2a-rpl10a2xHA)^530^*^531^*; Tg(huc:gal4)* double transgenic animal. **(E)** Schematic diagram illustrating the workflow for TRAP purification of RNAs from RiboTag Reporter zebrafish crossed to Gal4 driver lines. **(F)** Quantitative RT-PCR measurement of the relative expression of the endothelial-specific *kdrl* gene in samples prepared from TRAP purified RNA from *Tg(uas:egfp-2a-rpl10a2xHA)^530^*^531^*; Tg(fli1a:gal4ff)^ubs^*^4^ double-transgenic animals compared to RNA prepared whole embryo lysates from the same animals. **(G)** Quantitative RT-PCR measurement of the relative expression of the neural-specific *snap25* gene in samples prepared from TRAP purified RNA from *Tg(uas:egfp-2a-rpl10a2xHA)^530^*^531^*;Tg(huc:gal4)* double-transgenic animals compared to RNA prepared whole embryo lysates from the same animals.

## DISCUSSION

In this study, we introduce a novel AngioTag transgenic line for profiling global gene expression in zebrafish vascular endothelial cells in their undisturbed endogenous environment. We have performed TRAP-RNAseq profiling of the endothelial translatome of 24hpf AngioTag larvae, showing that TRAP-RNAseq results in superior endothelial gene enrichment compared to RNAseq on FACS-isolated endothelial cells. We have used AngioTag TRAP-RNAseq to document vascular enrichment of a number of annotated genes that were not previously known to be expressed in the endothelium, and to identify several novel unannotated vascular genes. We have also profiled the endothelial gene expression of several different adult organs, uncovering genes shared by the endothelium of all organs and genes unique to the vasculature of specific organs. Finally, to make TRAP-RNAseq methodologies accessible to all researchers, not just those studying the vascular endothelium, we have generated a UAS-RiboTag line that can be used to profile any cell and tissue type for which a Gal4 driver line is available.

Classically, translatomes have been prepared for profiling using sucrose density gradients, collecting RNA from the polysome fraction^40^. Profiling the translatome of a particular tissue type using this method requires either dissection or FACS sorting. More recently, the TRAP method has been used to profile the translatomes of specific tissues or cell types by using mice with a floxed conditional ‘tagged’ allele of the Rpl22 ribosomal protein expressed after crossing the mice to cell- or tissue-specific Cre driver lines^12^. RiboTag mice have been used to assay gene expression in hypothalamic neurons through TaqMan assays^41^, Sertoli and Leydig cells of the testes through microarrays^42^, proneural gliomas through ribosomal footprinting on translatome RNA^43^, factor VII expression in endothelial cells through qPCR^44^, and endothelial gene expression during homeostasis and inflammation through RNA-sequencing^45^. Other groups have utilized the TRAP method to perform RNAseq on specific cell populations in the mouse brain or kidney, with all but one group amplifying their RNA prior to sequencing^14,46–48^. Tissue specific RiboTag zebrafish have also been generated driving expression of GFP fused to rpl10a with tissue-specific promoters and then utilizing anti-GFP antibodies to immunoprecipitate tissue specific translatomes. Translatomes prepared in this way have been used to assay gene expression in the heart using microarrays^49^ and in melanocytes using qPCR^50^. A binary zebrafish RiboTag system has also been generated in which transgenic zebrafish carrying Avi-tagged Rpl10a are crossed to tissue specific BirA lines to biotinylate Rpl10a in BirA-expressing tissues. This binary biotinylation method was used to perform microarray analysis on zebrafish skeletal muscle mRNAs^51^. Notably, the two zebrafish studies noted above using microarrays also amplified their collected translatome RNA prior to further analysis. In contrast, our larval TRAP-RNAseq procedures were performed without any amplification of collected RNA.

Our whole embryo translatome TRAP-RNAseq results provide a valuable dataset for uncovering new insights into sequence features associated with more- or less-highly translated genes. Interestingly, our data on codon usage associated with highly translated genes shows strong parallels with previously published data examining codon usage in the developing zebrafish^33^. In that study, the authors examined codon usage in the 469 most highly transcribed genes in the zebrafish genome (as determined from previously published RNAseq data), identifying codons preferentially associated with highly transcribed genes. These authors went on to show that by replacing the native codons with codons preferred in “more transcribed” genes they were able to increase protein accumulation, providing experimental support for the idea that zebrafish use “optimal” codons to increase translation of their already highly transcribed genes. Our data reveals that many of the same codons enriched in the highly transcribed gene set of Horstick et al. are also preferentially enriched in the most-highly translated genes in our whole embryo translatome dataset (**Fig. 3C**), providing direct experimental evidence for the idea that highly transcribed genes are also “translationally optimized.”

Comparison between our TRAP-RNAseq and FACS-RNAseq endothelial datasets (**Fig. 4**) suggests that TRAP-RNAseq provides more effective enrichment of endothelial genes. Of the top twenty enriched GO terms in each dataset, there were more than twice as many vascular-related biological process GO terms in the TRAP-RNAseq dataset compared to the FACS-RNAseq dataset (**Supplemental Table 2**). Interestingly, principal component analysis of our RNAseq data (**Figure 2D**) also shows that the three AngioTag TRAP-RNAseq replicates (red spheres) are more highly clustered with one another and with the other non-FACS RNAseq datasets (light blue, dark blue, and yellow spheres) than the RNAseq data from either FACS-sorted endothelial cells (green spheres) or from dissociated but unsorted total embryonic cells (purple spheres), which are much less clustered, either with their own replicates or with the other RNAseq datasets. These results suggest that technical aspects of the FACS process, such as embryonic dissociation and prolonged incubation of separated cells prior to collection of mRNA (samples 3 and 4 in **Figure 2C**), introduce significant changes in gene expression compared to samples in which cells are rapidly disrupted prior to either immediate collection of mRNA (samples 1 and 5 in **Figure 2C**) or TRAP-RNAseq (samples 2 and 6 in **Figure 2C**). Using our TRAP-RNAseq methods, we were able to identify previously uncharacterized endothelial genes. These include both annotated genes not previously shown to have endothelial-enriched expression (**Figure 5A-F**) and unannotated genes (**Figure 5G-P**), some of which we have also shown are required for normal vascular development (**Figure 6**). These results further reinforce the idea that the AngioTag TRAP-RNAseq approach can provide unique insights compared to FACS sorting of dissociated endothelial cells.

We also show that AngioTag TRAP-RNAseq provides an effective method for querying and comparing the endothelial translatomes of different adult organs. Comparing the endothelial translatome of each organ to its own tissue lysate reveals shared endothelial signatures across all organs (**Figure 7B**). Pairwise comparisons between the endothelial translatomes of each organ also reveals unique, organ-specific endothelial profiles (**Figure 7I**), uncovering diversity among the vascular beds of different organs that likely relect functional differences between the vessels found in different tissues. These findings further reinforce the importance of profiling the vasculature within its native, *in vivo* environment since it is evident that tissue location plays an important role in vascular gene expression.

Finally, the new *Tg(uas:egfp-2a-rpl10a2xHA)^530^*^531^ UAS:RiboTag line that we have generated in the course of this work makes it possible to apply the TRAP-RNAseq methods we used to profile the endothelium to any cell or tissue type in the zebrafish for which a Gal4 driver line is available (**Figure 8**). A very large number of transgenic Gal4 driver lines have been generated in the zebrafish with expression in a wide assortment of different cells and tissues, including many lines derived from enhancer trap screens^52–55^. The availability of these lines should allow the wide application of our TRAP-RNAseq methods.

Together, the new tools and methods we have developed for transgene-assisted TRAP-RNAseq provide a valuable resource for cell- and tissue-specific in situ expression profiling in developing and adult zebrafish. By combining these new TRAP-RNAseq profiling tools with other genetic and experimental tools available in the zebrafish, we can study changes in gene expression due to gene mutations, disease states, or during regeneration in numerous different cell types. These new tools and methods greatly enhance our capability to analyze changes in translational landscapes within cells in their local microenvironments.

## ACKNOWLEDGEMENTS

The authors would like to thank members of the Weinstein laboratory for their critical comments on this manuscript, Harold Burgess and Jennifer Sinclair for the huC:gal4 and xa210:gal4 fish, andTianwei Li at the NICHD MGC sequencing core for performing the library construction and RNA sequencing.

