## Supplemental Data for "Profiling the endothelial translatome *in vivo* using ‘AngioTag’ zebrafish"

Supplemental Table 1: Readcounts for larval RNAseq analysis

| Sample | Total mapped reads | Uniquely mapped reads | % uniquely mapped reads | Reads mapped to multiple loci | % reads mapped to multiple loci |
|---|---|---|---|---|---|
| 1-1 RiboTag Whole Lysate | 49956134 | 42588557 | 85.25% | 2969994 | 5.95% |
| 1-2 RiboTag Whole Lysate | 69239038 | 59221672 | 85.53% | 3076200 | 4.44% |
| 1-3 RiboTag Whole Lysate | 47632455 | 41583642 | 87.30% | 2029387 | 4.26% |
| 2-1 RiboTag TRAP | 50661654 | 34986862 | 69.06% | 3703631 | 7.31% |
| 2-2 RiboTag TRAP | 56252261 | 43023522 | 76.48% | 3810600 | 6.77% |
| 2-3 RiboTag TRAP | 45403187 | 36331265 | 80.02% | 2344794 | 5.16% |
| 3-1 AngioTag FACS Total Cells | 55085749 | 51341850 | 93.20% | 1651529 | 3.00% |
| 3-2 AngioTag FACS Total Cells | 53707170 | 49094355 | 91.41% | 1495389 | 2.78% |
| 3-2 AngioTag FACS Total Cells | 58235304 | 51189179 | 87.90% | 2147287 | 3.69% |
| 4-1 AngioTag FACS ECs | 56759660 | 50662241 | 89.26% | 1443422 | 2.54% |
| 4-2 AngioTag FACS ECs | 55032277 | 31439590 | 57.13% | 20211551 | 36.73% |
| 4-3 AngioTag FACS ECs | 52123159 | 45630183 | 87.54% | 2572154 | 4.93% |
| 5-1 AngioTag Whole Lysate | 46722553 | 40175213 | 85.99% | 1951949 | 4.18% |
| 5-2 AngioTag Whole Lysate | 57354200 | 53786329 | 93.78% | 1898238 | 3.31% |
| 5-3 AngioTag Whole Lysate | 53618024 | 50073486 | 93.39% | 1900801 | 3.55% |
| 6-1 AngioTag TRAP | 49463459 | 26997399 | 54.58% | 7640140 | 15.45% |
| 6-2 AngioTag TRAP | 58760419 | 54794280 | 93.25% | 2027329 | 3.45% |
| 6-3 AngioTag TRAP | 55289491 | 51098279 | 92.42% | 2198844 | 3.98% |

Supplemental Table 2: Shared GO terms between FACs and AngioTag

| GO Biological Process | FACs fold enrichment | AngioTag fold enrichment | FACs p -value | AngioTag p-value |
|---|---|---|---|---|
| endothelial cell migration (GO:0043542) | 5.44 | 7.14 | 1.10E-02 | 6.73E-03 |
| lymph vessel development (GO:0001945) | 3.66 | 5.13 | 1.27E-02 | 6.01E-04 |
| blood vessel morphogenesis (GO:0048514) | 3.07 | 4.14 | 1.21E-14 | 6.55E-19 |
| blood vessel development (GO:0001568) | 3.05 | 3.94 | 6.72E-17 | 8.66E-20 |
| vasculature development (GO:0001944) | 2.94 | 3.85 | 3.44E-18 | 2.85E-22 |
| angiogenesis (GO:0001525) | 2.92 | 3.99 | 2.92E-09 | 7.04E-13 |

Supplemental Table 3: Top 10 GO terms for skin AngioTag

| Skin AngioTag: GO Biological Process | Number of genes in set | Mean log2fold change | Adjusted p-value |
|---|---|---|---|
| angiogenesis (GO:0001525) | 129 | 1.80 | 2.33E-08 |
| actin filament organization (GO:0007015) | 127 | 1.73 | 6.32E-09 |
| blood vessel morphogenesis (GO:0048514) | 163 | 1.72 | 6.72E-12 |
| small GTPase mediated signal transduction (GO:0007264) | 129 | 1.58 | 6.72E-12 |
| regulation of anatomical structure morphogenesis (GO:0022603) | 136 | 1.58 | 1.31E-09 |
| regulation of small GTPase mediated signal transduction (GO:0051056) | 61 | 1.53 | 1.25E-06 |
| positive regulation of transcription by RNA polymerase II (GO:0045944) | 118 | 1.50 | 1.24E-07 |
| regulation of GTPase activity (GO:0043087) | 87 | 1.47 | 1.61E-09 |
| chromatin organization (GO:0006325) | 116 | 1.41 | 2.33E-08 |
| RNA splicing (GO:0008380) | 112 | 1.31 | 1.32E-06 |

Supplemental Table 4: Top 10 GO terms for muscle AngioTag

| **Muscle AngioTag: GO Biological Process** | **Number of genes in set** | **Mean log2fold change** | **Adjusted p-value** |
|---|---|---|---|
| angiogenesis (GO:0001525) | 153 | 2.41 | 4.62E-20 |
| blood vessel morphogenesis (GO:0048514) | 183 | 2.33 | 1.47E-22 |
| chromatin organization (GO:0006325) | 127 | 1.58 | 1.13E-14 |
| histone modification (GO:0016570) | 102 | 1.51 | 4.58E-13 |
| regulation of mRNA metabolic process (GO:1903311) | 78 | 1.50 | 3.00E-12 |
| RNA splicing (GO:0008380) | 143 | 1.42 | 8.26E-21 |
| mRNA processing (GO:0006397) | 162 | 1.42 | 2.29E-20 |
| mRNA splicing, via spliceosome (GO:0000398) | 115 | 1.40 | 1.64E-19 |
| RNA splicing, via transesterification reactions (GO:0000375) | 115 | 1.40 | 1.64E-19 |
| RNA splicing via transeterfication reaction with bluged adenosine as nucleophile (GO:0000377) | 115 | 1.40 | 1.64E-19 |

Supplemental Table 5: Top 10 GO terms for liver AngioTag

| Liver AngioTag: GO Biological Process | Number of genes in set | Mean log2fold change | Adjusted p-value |
|---|---|---|---|
| lymphangiogenesis (GO:0001946) | 25 | 3.69 | 9.73E-09 |
| lymph vessel morphogenesis (GO:0036303) | 26 | 3.67 | 1.09E-89 |
| lymph vessel development (GO:0001945) | 35 | 3.36 | 1.07E-10 |
| sprouting angiogenesis (GO:0002040) | 49 | 3.20 | 1.81E-08 |
| angiogenesis (GO:0001525) | 124 | 3.00 | 4.66E-22 |
| blood vessel morphogenesis (GO:0048514) | 148 | 2.96 | 2.48E-25 |
| regulation of anatomical structure morphogenesis (GO:0022603) | 97 | 2.72 | 3.31E-09 |
| ameboidal-type cell migration (GO:0001667) | 79 | 2.65 | 1.94E-09 |
| actin filament organization (GO:0007015) | 93 | 2.51 | 2.89E-09 |
| small GTPase mediated signal transduction (GO:0007264) | 98 | 2.50 | 3.06E-13 |

Supplemental Table 6: Top 10 GO terms for heart AngioTag

| **Heart AngioTag: GO Biological Process** | **Number of genes in set** | **Mean log2fold change** | **Adjusted p-value** |
|---|---|---|---|
| actin filament organization (GO:0007015) | 86 | 1.69 | 1.76-09 |
| angiogenesis (GO:0001525) | 101 | 1.68 | 1.41E-14 |
| blood vessel morphogenesis (GO:0048514) | 120 | 1.64 | 2.14E-16 |
| stem cell differentiation (GO:0048863) | 70 | 1.59 | 5.32E-08 |
| embryonic organ morphogenesis (GO:0048562) | 103 | 1.58 | 2.67E-09 |
| regulation of multicellular organismal development (GO:2000026) | 95 | 1.58 | 4.50E-08 |
| small GTPase mediated signal transduction (GO:0007264) | 78 | 1.56 | 4.50E-08 |
| regulation of anatomical structure morphogenesis (GO:0022603) | 91 | 1.53 | 9.55E-10 |
| skeletal system development (GO:0001501) | 93 | 1.53 | 1.76E-09 |
| chromatin organization (GO:0006325) | 84 | 1.52 | 9.32E-11 |

Supplemental Table 7: Top 10 GO terms for brain AngioTag

| Brain AngioTag: GO Biological Process | Number of genes in set | Mean log2fold change | Adjusted p-value |
|---|---|---|---|
| sprouting angiogenesis (GO:0002040) | 47 | 3.28 | 3.20-05 |
| angiogenesis (GO:0001525) | 113 | 3.23 | 1.18E-11 |
| blood vessel morphogenesis (GO:0048514) | 137 | 3.18 | 5.34E-14 |
| regulation of anatomical structure morphogenesis (GO:0022603) | 95 | 2.49 | 3.20E-05 |
| regulation of cells shape (GO:0008360) | 34 | 2.47 | 8.55E-06 |
| Ras protein signal transduction (GO:0007265) | 70 | 2.45 | 7.48E-07 |
| small GTPase mediated signal transduction (GO:0007264) | 95 | 2.35 | 4.27E-08 |
| regulation of hydrolase activity (GO:0051336) | 99 | 2.34 | 3.20E-05 |
| regulation of cell morphogenesis (GO:0022604) | 41 | 2.31 | 9.83E-07 |
| regulation of cysteine-type endopeptidase activity involved in apoptotic process (GO:0043281) | 23 | 2.28 | 3.20E-05 |
